# YeastIT: Reducing mutational bias for in vivo directed evolution using a novel yeast mutator strain based on dual adenine-/cytosine-targeting and error-prone DNA repair

**DOI:** 10.1101/2023.11.20.567881

**Authors:** Marta Napiorkowska, Katrin Fischer, Matthew Penner, Philipp Knyphausen, F. Hollfelder

## Abstract

Engineering proteins with new functions and properties often requires navigating large sequence spaces through rounds of iterative improvement. However, a disparity exists between the gradual pace of natural long-term evolution and a typical laboratory evolution workflow that relies on enriching functional variants from highly diverse in vitro generated libraries through very few screening rounds. Laboratory experiments often eschew presumed natural strategies such as neutral/non-adaptive and multi-phase evolution trajectories, and therefore mutagenesis technologies suitable for long ‘nature-like’ timescales are needed. Here, we introduce YeastIT, a novel in vivo mutagenesis tool for protein engineering that leverages an *S. cerevisiae* strain engineered to exhibit mutagenic activity directed to the gene of interest, allowing its continuous diversification. Mutagenesis is achieved by generating DNA damage through nucleoside deamination, followed by introduction of mutations by harnessing the process of error-prone DNA translesion synthesis. By eliminating the transformation step, YeastIT allows multiple rounds of screening or selection without interruptions for library diversification, thereby enabling long-term and continuous evolution campaigns. Our characterization of the mutational spectrum and frequency of the YeastIT-generated libraries, and its comparison to other methods (error-prone PCR, PACE, MutaT7, eMutaT7, OrthoRep, TRIDENT, EvolVR) demonstrates comparable mutation rates combined with a significant reduction in mutagenic bias relative to most of the alternatives. To validate YeastIT, we carried out directed evolution of a DARPin binding protein to achieve a 15-fold improved affinity. YeastIT thus provides a tool for exploring different evolutionary trajectories which overcomes previous limitations of variant availability (due to bias and low mutation rates) and emulates the way proteins emerge in Nature.

## Introduction

Natural protein evolution is characterised by trajectories extending over many rounds and millions of years, as functional solutions can be rare and challenging to discover in vast sequence spaces. Cells provide mechanisms for continuous (albeit slow) randomisation by virtue of being evolutionary units that combine genotype and phenotype in one compartment. By contrast, directed evolution of proteins in the laboratory is characterised by much shorter trajectories, partially because of experimental considerations to accommodate workflows that alternate between in vitro and in vivo steps in each round in a reasonable timeframe. Typically, in vitro DNA diversification (e.g., by error-prone PCR, site-directed mutagenesis, DNA shuffling) is followed by the introduction of variants into living hosts for functional high-throughput screening, creating a substantial experimental burden.

In vivo mutagenesis presents an attractive alternative to in vitro genetic diversification and practically simplifies directed evolution campaigns^1^. By enabling a continuous workflow of mutagenesis and selection, several significant advantages can be realised: (i) bypassing iterative rounds of in vitro DNA manipulation, transformation into host cells, and in vivo phenotypic screening accelerates the speed at which each round can be completed and saves costs in labour and reagents; (ii) elimination of repetitive transformation steps enables generation of larger DNA libraries as their size is often restricted by transformation efficiency when generated in vitro; and (iii) in the instances where the function of interest can be efficiently coupled to the selective pressure on the host, the coverage of larger regions of sequence space becomes possible with minimal experimental effort, thereby unlocking access to phenotypes that require very long evolutionary pathways.

Methods for gene-targeted mutagenesis in vivo (**Table 1**) can be broadly divided into two categories based on their underlying molecular mechanism. One approach relies on introducing mutations into a plasmid or genomic locus containing the target gene using an error-prone DNA^2–5^ or RNA polymerase^6^ variant that is orthogonal to the host (i.e. does not modify its native genome). An alternative approach uses nucleoside deaminases fused to a gene-targeting partner such as RNA polymerase^7–10^ (enabling whole-gene mutagenesis) or a deactivated Cas endonuclease^11,12^ (enabling targeted DNA base editing). Both mechanisms have been demonstrated in bacteria^2,3,7,8^, yeast^4,5,9^, and mammalian cells^6,10^.

**Table 1.** Overview of existing technologies for in vivo mutagenesis.

| Method | Host organism | Molecular mechanism | Mutation rate<sup>a</sup><br>(mut/kb/day) | Mutation spectrum | Mutagenic window | Validation in directed evolution |
| --- | --- | --- | --- | --- | --- | --- |
| MutaT7<sup>7</sup> | *E. coli* | T7 RNA polymerase fused to cytidine deaminase | 0.34 | GC>AT only | ~1 kb | Drug resistance |
| eMutaT7<sup>8</sup> | *E. coli* | T7 RNA polymerase fused to cytidine deaminase | 3.7 | GC>AT only | ~1 kb | Drug resistance |
| EvoIVR<sup>2</sup> | *E. coli* | nCas9 fused to error-prone polymerase Poll | 0.09 | All types | 50 nt/gRNA<sup>b</sup> | Drug resistance, enzyme activity |
| PACE<sup>3</sup> | *E. coli* | GOI linked to phage fitness; non-targeted error-prone Poll | 1.25 | All types | >2 kb | Enzyme activity, transcription factor affinity |
| CdA-RNAPT7<sup>18</sup> | *Pseudomonas putida* | T7 RNA polymerase fused to cytidine deaminase | n/d | GC>AT only | ~1 kb | Drug resistance |
| BacORep<sup>19</sup> | *Bacillus thuringiensis* | Error-prone GIL16 DNAP specific for linear plasmid | 0.45 | All types<sup>c</sup> | >2 kb | Promoter strength |
| ICE<sup>5</sup> | *S. cerevisiae*; *Kluyveromyces lactis* | error-prone self-replication of Ty1 retrotransposon | 0.05 | All types<sup>c</sup> | >2 kb | Drug resistance, metabolic engineering |
| OrthoRep<sup>4</sup> | *S. cerevisiae* | error-prone TP DNAP1 specific for p1 plasmid | 0.6 | >90% transitions | >2 kb | Drug resistance, enzyme activity, binder affinity |
| TRIDENT<sup>9</sup> | *S. cerevisiae* | T7 RNA polymerase fused to cytidine and adenine deaminases | 0.7 | >97% transitions<sup>d</sup> | ~1 kb | Drug resistance |
| TRACE<sup>10</sup> | HEK cells | T7 RNA polymerase fused to cytidine deaminase | 0.7 | GC>AT only | >2 kb | Drug resistance |
| VEGAS<sup>6</sup> | HEK cells | GOI linked to Sindbis virus fitness; virus-specific RNA replicase | 2.4 | All types | ~1 kb | Transcription factor affinity |

<sup>a</sup> Mutation rate of best-performing construct reported.
<sup>b</sup> Simultaneous use of 2 gRNAs was demonstrated for plasmid mutagenesis. For genome mutagenesis, adjacent gRNAs must be located on the same strand and confer lethal phenotype otherwise.<sup>2</sup>
<sup>c</sup> Mutational spectra of BacORep and ICE, as reported by the authors based on Illumina sequencing data, cannot be considered accurate since the observed frequencies are comparable to the sequencing error rate. In the case of BacORep, this dominance of sequencing errors is further evidenced by the lack of differences in mutational spectra between the mutagenesis sample and the negative (non-passaged) control and was confirmed by our reanalysis. We were unable to conduct a similar reanalysis for ICE due to the lack of sufficient publicly available data.
<sup>d</sup> The authors report 97 % transitions. In this study we adjusted the analysis to cover the full length of the gene without altering other parameters of the analysis and obtained a transition rate of 99%.

Laboratory yeast strains such as *Saccharomyces cerevisiae* are a particularly attractive platform for directed evolutions as they combine the advantages of bacterial systems (fast replication rates, low-cost and straightforward culturing, availability of a well-established toolbox for genetic manipulation)^13^ with those of mammalian systems (expression of eukaryotic proteins, ability to perform post-translational modifications, high secretory capacity)^14^. Several in vivo gene diversification methods have been demonstrated in yeast. Crook et al. developed in vivo continuous evolution (ICE), which involves introducing a gene of interest as a cargo on the engineered version of the retrotransposon Ty1 and subjecting it to cycles of error-prone reverse transcription and subsequent genomic reintegration^5^. ICE was successfully applied in *S. cerevisiae* to evolve the transcriptional regulator Spt15 to confer improved tolerance to 1-butanol, and the xylose metabolic pathway to increase the rate of consumption of xylose. However, ICE exhibits a relatively low mutation rate (**Table 1**; 0.05 mutations/kb/day, approximately 10,000-fold higher than natural genomic error rates^1^), leading to slow adaptive walks, and was reported to suffer from undesired reintegration of mutants into alternative loci in the genome, which increases the copy number of the gene of interest, thus impairing selection efficiency^5^.

An alternative method, called OrthoRep, was reported by Ravikumar et al. and relies on the genome-orthogonal error-prone polymerase TP-DNA1 which replicates the cytoplasmic plasmid p1, introducing mutations to a p1-encoded gene of interest^4^. This method exhibits a 12-fold improved mutation rate relative to ICE but suffers from a substantial bias in mutational diversity (more than 90% of mutations are transition mutations, i.e. A/G or C/T interchanges). Random introduction of mutations throughout the plasmid precludes precise targeting to the gene of interest, effectively wasting mutational load on the genetic elements whose modification is undesirable. Despite these limitations, OrthoRep was successfully applied to improve the activity of the tryptophan synthase TrpB^15^ and the binding affinity of a nanobody against SARS-CoV-2 RBD^16^.

Fusion of nucleoside deaminases with RNA polymerases has been implemented in yeast by Cravens et al. in TRIDENT, where mutations are introduced by the activity of an engineered adenosine deaminase variant and sea lamprey cytidine deaminase aided by two co-localized DNA repair factors^9^. Cytidine deaminase catalyses the deamination of deoxycytidine to deoxyuridine (C>U), while adenosine deaminase catalyses the deamination of deoxyadenosine to deoxyinosine (A>I). During DNA replication, dATP and dCTP are inserted opposite deoxyuridine and deoxyinosine, respectively, leading to G>A and T>C mutations. Each deaminase is fused to a separate T7 RNA polymerase (T7RNAP) partner, which targets its activity specifically towards the gene of interest flanked by a T7 promoter and a T7 terminator. The system shows comparable mutational rates to OrthoRep but even higher bias (> 97%) towards transition mutations. The methods that employ bacteria as host organisms show similar disadvantages of low mutational rate (EvolvR^2^) and limited spectrum (deaminase-based systems such as MutaT7 and its derivatives^7,8^), or suffer from other limitations such as complicated experimental setups and the requirement for coupling of function of interest to viral fitness (PACE^3^; see **Table 1** for extended comparison of mechanisms and parameters in existing methods).

We establish a novel method for gene mutagenesis called YeastIT (Yeast *In vivo* Targeted mutagenesis) that relies on the previously applied principle of fusing nucleoside deaminase and T7 RNA polymerase. We show that our design features a significantly more diversified mutational spectrum than TRIDENT and all other methods that exploit this mechanism, while showing comparable mutational rates. Furthermore, we perform a new comparative analysis of the mutational spectra achieved by YeastIT at both nucleotide and amino acid levels, in comparison to established random mutagenesis methods in yeast (OrthoRep, TRIDENT), *E. coli* (PACE, MutaT7, eMutaT7, EvolvR) and in vitro (error-prone PCR with Taq and with Mutazyme II). This comprehensive analysis highlights major differences in sequence space accessibility between different methods and provides a valuable reference for selecting the optimal mutagenesis strategy for a directed evolution campaign. Finally, we validate YeastIT by evolving an EGFP-binding designed ankyrin repeat protein^17^ (DARPin) for enhanced binding affinity and identify a variant with 15-fold improved binding, establishing the first directed evolution campaign involving deaminase-based hypermutators that extends beyond applications linked to a growth selection.

## Methods

### Strains, materials, and molecular biology techniques

The strains used in this study are listed in **Supplementary Table 1**, plasmids in **Supplementary Table 2**, primers in **Supplementary Table 3**. Proteins sequences are given in **Supplementary Table 4**, **Supplementary Table 5** and **Supplementary Figure 1**. *E. coli* NEB5-alpha (NEB#C2987) was used for the construction of all plasmids. *E. coli* BL21(DE3) (NEB#C2527) was used for protein overexpression. Restriction enzymes were obtained from ThermoFisher or NEB. All in vitro PCR reactions were performed using Q5 High-Fidelity 2X Master Mix (NEB). Yeast colony PCR reactions were done as previously reported^20^ using DreamTaq DNA polymerase (ThermoFisher). Gibson Assembly reactions were done using Gibson Assembly Master Mix (NEB). Oligonucleotides were obtained from Merck. Synthetic DNA fragments larger than 50 bp were obtained from Integrated DNA Technologies as gBlocks. For plasmid DNA purification, GeneJET Miniprep kit (ThermoFisher) and Zymoprep Yeast Plasmid Miniprep II kit (Zymo Research) were used for bacterial and yeast cultures, respectively. For the purification of PCR products, Zymo Clean&Concentrator Kit (Zymo Research) was used. Yeast transformations (plasmids and linear fragments for genomic integration) were done using the Frozen-EZ Yeast Transformation II Kit (Zymo Research). All other chemicals were obtained from Merck unless otherwise specified.

### Microbial cultivations

*E. coli* strains were grown in LB medium supplemented with 50 µg/mL kanamycin or 100 µg/mL ampicillin when required, on an orbital shaker (37°C, 200 rpm, 19 mm amplitude). For protein expression in *E. coli*, cultivation conditions were changed to 25°C, 300 rpm, 25 mm amplitude. *S. cerevisiae* strains were cultured in yeast extract/peptone/dextrose medium (YPD, 1% yeast extract, 2% peptone, 2% dextrose) when containing no plasmids and in synthetic defined (SD) media (0.67% yeast nitrogen base without amino acids [BD Difco], 2% dextrose) supplemented with the appropriate yeast synthetic drop-out medium supplement when plasmid maintenance was required (SDΔW-without tryptophan, SDΔL-without leucine, or SDΔLW-without leucine and tryptophan). For induction of the mutagenesis system, SD media were supplemented with doxycycline (0.25 µg/ml unless specified otherwise). For induction of mutagenesis target expression, media supplemented with 2% galactose instead of 2% dextrose was used (SGΔW, SGΔL, SGΔLW). All *S. cerevisiae* cultivations were done at 30°C in an orbital shaker (1,000 rpm, 1.5 mm amplitude). For cultivations on solid media, 2% agar was added.

### Construction of *S. cerevisiae* strains with genomic knockouts and integrations

To construct strain EBY100 Δ*ung1,* gene *ung1* was deleted by amplifying the KanMX cassette from the plasmid HO-poly-KanMX4-HO with primers F1 and R1 containing overhangs homologous to regions upstream and downstream of the *ung1* locus and transforming the linear purified product into chemically competent *S. cerevisiae* EBY100. Cells were recovered for 5 hours, plated on YPD-agar supplemented with 100 µg/ml G418 disulphate salt, and the successful deletion was confirmed by colony PCR. Strain EBY100 Δ*apn1* was constructed in the same way, but primers F2 and R2 were used instead. To construct strains EBY100 Δ*ung1 HO::EVOAPO1* and EBY100 Δ*apn1 HO::YEASTIT2*, plasmids HO-evoAPO1_T7RNAP-LEU-HO and HO-evoAPO1_evoTadA_T7RNAP-LEU-HO were digested with *HindIII* and *ScaI* and purified inserts were transformed into chemically competent *S. cerevisiae* EBY100 Δ*ung1* or *S. cerevisiae* EBY100 Δ*apn1.* Cells were recovered for 5 hours, plated on SDΔL-agar, and successful integration was confirmed by colony PCR.

### In vivo *she1* mutagenesis

To perform in vivo mutagenesis of *she1* for fluctuation assays and nanopore sequencing, strains engineered to exhibit the mutator phenotype were transformed with pGoi-she1 and plated on SD-agar plates. A single colony was used to inoculate 500 µL of SD medium in a deep well plate (#780270, Greiner Bio-One) and grown to stationary phase (approximately 24 hours). Subsequently, 10 µL of saturated culture was used to inoculate 500 µL of SD medium supplemented with doxycycline and grown for 24 hours. The process was repeated 2 more times for a total of 72 hours. For each genotype tested, 3 biological replicates were always performed. The generated libraries were then used for the She1 fluctuation assay or for the UMIC-seq analysis.

### She1 fluctuation assay

Following the in vivo mutagenesis process, the cell suspensions were centrifuged (1,000 × g, 5 min), washed once to remove the remaining glucose, and resuspended in equal volume of PBS. Cells were serially diluted (dilution ratio from 10^-1^ to 10^-8^) in PBS, and 10 µL of each dilution was spotted on SD-agar and SG-agar plates. SD-agar plates were incubated for 48 hours and SG-agar plates were incubated for 96 hours to yield colonies of comparable size. To estimate mutational frequencies, the number of survivals on SG-agar and SD-agar plates was counted and the ratio between the two was calculated. For all fluctuation assays, the survival ratios of three biological replicates are reported and error bars represent mean±standard deviation. To identify the mutations, plasmids were recovered from yeast, propagated in *E. coli* NEB5-alpha, and *she1* inserts were Sanger-sequenced (Sequencing facility, Department of Biochemistry, University of Cambridge).

### Nanopore library preparation and UMIC-seq

Libraries generated using in vivo *she1* mutagenesis protocol with YeastIT1 and YeastIT2 strains were used as the input for the preparation of a nanopore sequencing library. For each strain, 3 biological replicates were pooled together and used for plasmid isolation. The plasmid library was transformed into *E. coli* NEB5-alpha (∼10,000 transformants), colonies were scraped and used for plasmid isolation. The *she1* insert was amplified with forward primers containing 56 nt-long unique molecular identifier (UMI; F3/F4) and reverse primers containing 24 nt-long sample-specific barcode (R3/R4) for 4 cycles, and with primers F5 and R5 for 15 cycles. The PCR product was gel-purified, Gibson-assembled with *SacI-* and *NcoI*-digested, gel-purified pASK-IBA63b+, and transformed into *E. coli* NEB5-alpha. Plasmids were isolated from scraped colonies (∼8,000 for YeastIT1, ∼4,000 for YeastIT2), digested with *SacI* and *NcoI*, subjected to gel purification followed by AMPure XP beads purification (Beckman Coulter), mixed in equimolar concentration (57 fmol/sample), and used for sample preparation with the SQK-LSK110 sequencing library preparation kit (Oxford Nanopore Technologies). The prepared sample (50 fmol) was sequenced using MinION sequencer equipped with an R9.4.1 flow cell (ONT). Sequencing generated 8.84 million reads (16.11 Gb), which corresponded to >600-fold oversampling. Base calling was performed with Guppy (v. 4.3.4) using default parameters. NanoFilt (v. 2.8.0) was used to filter sequences based on length (1,750-2,000 bp) and read quality score (8). Filtered reads were demultiplexed based on barcode sequence (TCGATTCCGTTTGTAGTCGTCTGT and GAGTCTTGTGTCCCAGTTACCAGG), yielding 3.81 million reads for YeastIT1 and 2.04 million reads for YeastIT2, and clustered based on UMI sequence identity (alignment threshold of 70, cluster size threshold of 50; error rate below 0.01% expected at these parameters^21^), yielding 10,503 and 4,336 clusters with median sizes of 488 and 733 sequences for YeastIT1 and YeastIT2 libraries, respectively. Consensus sequences of each cluster were generated with medaka (v 1.2.2; default parameters).

### Yeast surface display, immunostaining, and flow cytometric analysis of DARPin 3G146

To induce surface display of DARPin 3G146, yeast cells were grown to stationary phase in SDΔLW or SDΔW, centrifuged (1,000 × g, 5 min), resuspended in equal volume of PBS, used to inoculate SGΔLW or SGΔW at 1:10 dilution, and grown for 24 hours. To detect DARPin expression level, cells were washed with PBS, diluted to OD of 0.1 – 0.3 and stained with THE^TM^ iFluor 647 HA Tag antibody (GenScript; 1:500 – 1:1,000 dilution of 0.5 mg/ml stock in 10 mg/ml BSA, PBS, pH 7.4) for 1 hour at room temperature on a roller shaker. To detect EGFP binding, cells were simultaneously stained with EGFP-His^6^ (117.5 μM stock in 10% glycerol, PBS, pH 7.4; concentration specified for each experiment). Subsequently, cells were diluted 2-fold with PBS and analysed with CytoFLEX S (Beckman Coulter; EGFP, λ^Ex^ 488 nm, λ^Em^ 525/40 nm; iFluor 647, λ^Ex^ 638 nm, λ^Em^ 660/10 nm).

### Library generation and flow cytometric sorting of yeast surface-displayed DARPin

To generate the DNA library of surface-displayed DARPin 3G146, YeastIT1 and YeastIT2 strains containing the plasmid pGoi-Aga2-3G146 were grown in SDΔLW and subjected to the same protocol as described for in vivo *she1* mutagenesis. Subsequently, cells were centrifuged (1,000 × g, 5 min), resuspended in an equal volume of PBS, and the display of DARPin variants was induced by transferring 40 µL of cell suspension to 460 µL of SGΔLW and growing for another 24 hours. Cells were centrifuged (1,000 × g, 5 min), washed once, resuspended in PBS to OD of 3.33, and stained with EGFP-His_6_ (90 nM in the first round, 30 nM in the second round) and iFluor 647 HA Tag antibody (1:1,000 dilution). Cells were diluted 2-fold with PBS, passed through a 10 µm cell strainer, and sorted with the FACSAria III system (BD Life Sciences). In the first round, a total of 1.9 million cells were analysed and 36 cells with the highest ratio of EGFP/iFluor 647 were sorted (**Supplementary Figure 2a**). Sorted cells were recovered on SDΔLW-agar plates, regrown colonies from the YeastIT1 and YeastIT2 sorting were pooled together and resubjected to the mutagenesis and sorting protocol. In the second round, a total of 1.4 million cells were analysed and 87 cells with the highest ratio of EGFP/iFluor 647 were sorted on SDΔLW-agar plates (**Supplementary Figure 2b**). Individual colonies were picked, used to inoculate liquid SDΔLW media, subjected to the yeast surface display, staining, and flow cytometric analysis protocol (OD of 0.3, 1:1,000 iFluor 647 HA Tag antibody, 30 nM EGFP-His_6_), and the EGFP signal of display-positive population was analysed to select the best-performing candidates for sequencing and further characterization.

### Binding affinity measurements of yeast-displayed DARPin variants

To determine the binding affinity of yeast-displayed DARPins, yeast cells were cultured, induced to display DARPin variants, stained according to equilibrium labelling protocol^22^ (culture OD of 0.1, 1:500 iFluor 647 HA Tag antibody, 1.6 nM – 1.6 μM EGFP-His_6_, 2 biological replicates), and subjected to flow cytometry at the same conditions as described above for 3G146. For each sample, the population was split into display-positive and display-negative fractions based on iFluor 647 signal and median EGFP fluorescence was determined for both. The median background fluorescence of the display-negative population at each EGFP-His_6_ concentration was subtracted from that of the display-positive one and the background-corrected median EGFP fluorescence values were used to calculate the apparent *K*_d_ values according to the formula 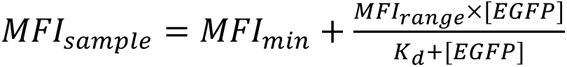as described before^23^.

### Overexpression and purification of DARPin variants

C-terminally His_6_-tagged DARPin variants were expressed in *E. coli* BL21(DE3) from plasmids pET-28a-3G146, pET-28a-3G146-2G4, and pET-28a-3G146-2H3 and subsequently purified by immobilized metal affinity chromatography (IMAC). A 2 mL LB preculture of each strain was grown overnight and used to inoculate 300 mL of TB at a dilution of 1:500. Upon reaching OD of 0.2, expression was induced with 200 μM IPTG. Cells were grown to stationary phase, harvested by centrifugation (4,000 × g, 30 min), and lysed by resuspending in 30 mL of wash buffer (50 mM Tris-HCl, pH 7.0, 300 mM NaCl, 20 mM imidazole, 10% glycerol) supplemented with 1x Bug Buster, 250 U of benzonase and 30 kU of rLysozyme and incubating 30 min on a roller shaker. Cell debris was removed by centrifugation (20,000 × g, 30 min) and the cell-free extract was loaded onto column packed with 2 mL Super Ni-NTA matrix (Protein Ark) pre-equilibrated with 3 column volumes (CV) of wash buffer. Column was washed with 5 CV of wash buffer and proteins were eluted with elution buffer (50 mM Tris-HCl pH 7.0, 300 mM NaCl, 250 mM imidazole, 10% glycerol) in 5 fractions of 1 mL. The fractions containing the protein were pooled and buffer was exchanged to PBS by concentrating 4 times on Amicon Ultra-15 Centrifugal filter (NMWL 10 kDA). The concentration was determined spectrophotometrically at a wavelength of 280 nm and using a molar extinction coefficient of 23,950 M^−1^ cm^−1^.

### Enzyme-linked immunosorbent assay (ELISA) of DARPin variants

Pierce™ Streptavidin Coated High Capacity plates (ThermoFisher) were coated with biotin-conjugated mouse anti-HA monoclonal antibody (10 μg ml^−1^; 2-2.2.14, ThermoFisher) in ELISA buffer (25 mM Tris-HCl, 150 mM NaCl, 0.1% BSA, 0.05% Tween 20, pH 7.2) at 4 °C overnight and washed three times with 200 μL ELISA buffer. DARPin variants (50 nM) were incubated with serially diluted EGFP-His_6_ (1.6 nM – 1.6 μM) in ELISA buffer for 2 hours at room temperature and 30 μL of each dilution was added to anti-HA-coated plates and incubated for another 1 hour. Plates were washed three times with 200 μL of ELISA buffer before adding rabbit anti-GFP antibody (1:10,000) in 100 μL of ELISA buffer, incubating 30 min at room temperature, repeating the washing steps, adding HRP-conjugated anti-rabbit secondary antibody (1:5,000) in 100 μL of ELISA buffer, incubating 30 min at room temperature, and repeating the washing steps. 50 μL of 1xTMB solution (ThermoFisher) were added to each well and incubated for 10 minutes. Reactions were terminated by adding 50 μL of 1M orthophosphoric acid and absorbance (450 nm) was measured with SpectraMax iD5 microplate reader (Molecular Devices).

### Biolayer Interferometry (BLI) of DARPin variants

Biotinylated EGFP was prepared using the ImmunoProbe™ Biotinylation Kit according to the manufacturer’s instructions using 4.2 mg/mL EGFP in PBS and a BAC-sulfoNHS:EGFP ratio of 2.5:1. After elution, biotinylated EGFP was concentrated using an Amicon Ultra-15 Centrifugal filter (NMWL 10 kDA). BLI experiments were performed using a ForteBio Octet RED96 system (Sartorius). Streptavidin-coated biosensors were hydrated in kinetics buffer (PBS, 0.1 mg/mL BSA, 0.01% Tween 20) for 10 minutes and biotinylated EGFP (65 μM) was captured for 120 s followed by baseline equilibration. Association of serially diluted DARPin variants (1,000 nM – 4.11 nM) was measured for 180 s followed by measuring dissociation in kinetics buffer for 180 s. Signals from control wells without DARPin and control biosensors without EGFP were subtracted before performing global data fitting to a 1:1 kinetic binding model to determine the association and dissociation rates.

### Data analysis

Flow cytometry data were analysed using FlowJo (v. 10.7.2, BD Biosciences). BLI data were analysed using Octet DataAnalysis (v. 11.0.0.4). Protein structures were visualized using PyMOL (v. 2.5.5). UMIC-seq analysis was done using Python (v. 3.7.1) with scripts adopted from https://github.com/fhlab/UMIC-seq and using R (v. 4.2.1) with maftools (v. 2.14.0). For a comparative analysis of mutagenesis methods, data were retrieved from original papers or reanalysed as specified in **Supplementary Table 7**. Analysis of TRIDENT data was performed with Python scripts adopted from the original publication^9^. Amino acid substitution matrices and force-directed graphs were generated using custom Python scripts with NetworkX (v. 2.4) package. On-yeast apparent binding affinities of DARPin variants were quantified using custom scripts in MATLAB (v. R2019b).

## Results

### Design of an in vivo mutagenesis system

Among the wide range of in vivo mutagenesis approaches **(Figure 1a**), yeast-based systems^4,5,9^ stand out for their robust and versatile protein expression capabilities and deaminase-T7RNAP fusions^7–10^ are attractive due to their precise targeting of mutations towards specific genes and compatibility with both growth selection and screening set-ups. However, none of the existing systems have fully met the critical requirement for effective in vivo mutagenesis: achieving a sufficiently high mutational rate to generate diversity comparable to in vitro mutagenesis methods, while simultaneously exhibiting minimal bias, particularly towards transition mutations. Building on the deaminase-T7RNAP concept, we developed an improved system that overcomes the limitations of the current methods with regard to mutational bias.

**Figure 1.**
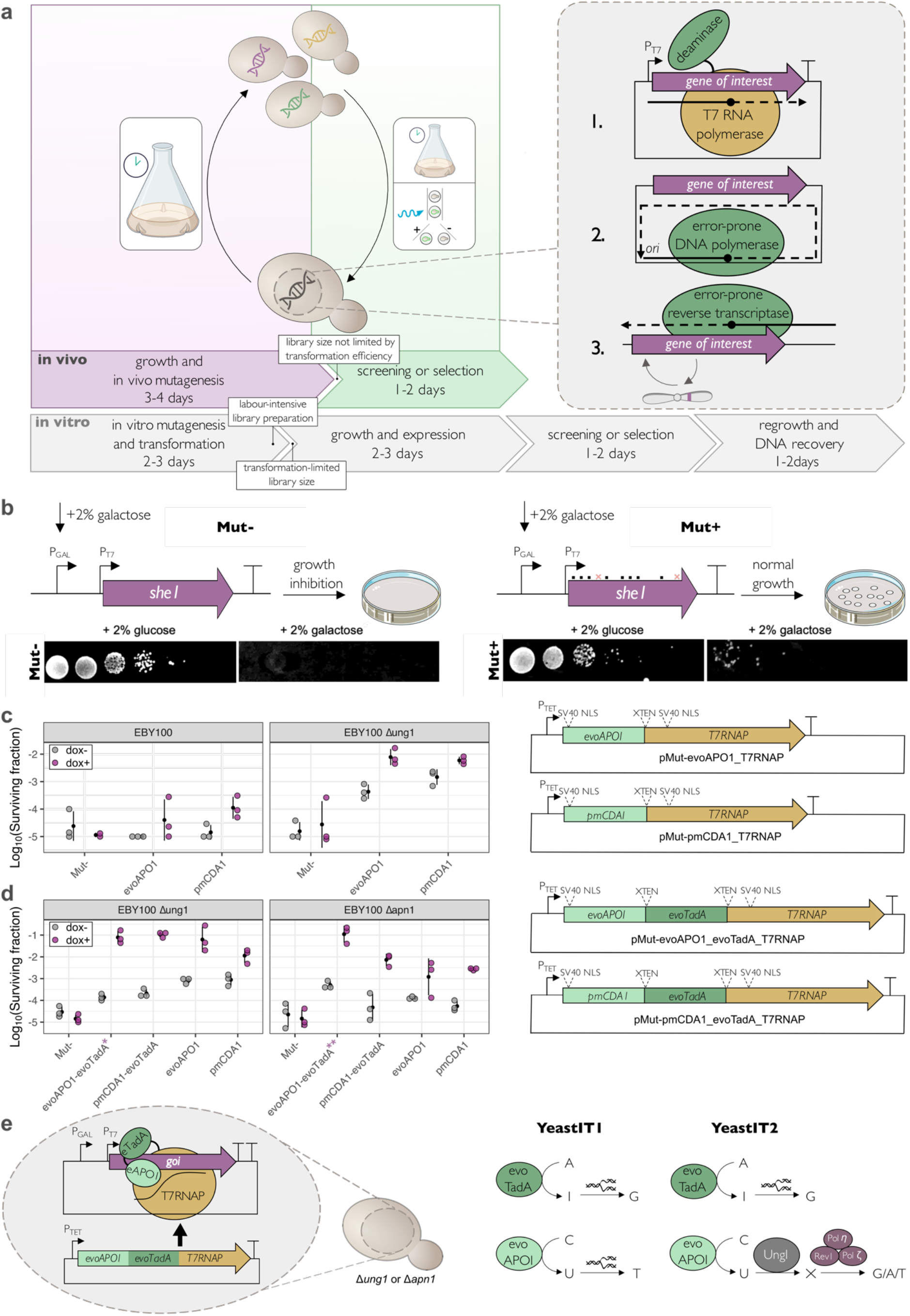
Development of the in vivo mutagenesis system. (**a**) Schematic overview of directed evolution workflows utilizing different mutagenesis strategies. In vivo mutagenesis can be accomplished with the use of T7 RNA polymerase-fused nucleoside deaminases (**1**), error-prone DNA-dependent polymerases (**2**) or RNA-dependent polymerases (**3**). Mutagenic protein domains are shown in green and accessory protein domains are shown in yellow. A timeline of typical directed evolution workflow utilizing in vitro mutagenesis is shown for comparison. (**b**) Schematic diagram of the fluctuation assay used to characterize the mutagenic performance of individual variants. When its expression is induced with galactose, *she1* confers a lethal phenotype to yeast. If galactose is not present, the growth rate is unaffected. Better performing mutagenesis systems lead to a higher proportion of cells containing deleterious mutations in *she1*, which improves the survival rate in galactose-containing media. (**c**) Mutagenic performance (left) and genetic architecture (right) of two mutator variants consisting of cytidine deaminase fused to T7RNAP in EBY100 or EBY100 Δ*ung1* genetic background. Both deaminase variants show similar deamination efficiency, while deletion of *ung1* leads to a 100-fold improvement in mutation efficiency. A weak mutagenic phenotype is also observed when the expression of mutator protein is not induced with doxycycline. XTEN, linker. SV40 NLS, nuclear localization signal. (**d**) Mutagenic performance (left) and genetic architecture (right, only variants not shown previously in **c**) of four mutator proteins consisting of adenosine and cytidine deaminases fused to T7RNAP in EBY100 Δ*ung1* or EBY100 Δ*apn1* genetic background. YeastIT1(Δ*ung1*) and YeastIT2(Δ*apn1*) strains are indicated with purple stars. (**e**) Schematic overview of final YeastIT1 and YeastIT2 mutagenesis strategies. In both strains, the mutator complex consists of adenosine deaminase evoTadA, cytidine deaminase evoAPO1, and T7RNAP. EvoTadA enables introduction of targeted A>G (T>C) mutations, while epoABO1 enables introduction of targeted C>T (G>A) mutations. T7RNAP targets the mutagenic activity towards the gene of interest that is located downstream of the T7 promoter sequence. In YeastIT2, mutations are detected and repaired by the native DNA repair machinery of the host in an error-prone fashion, thereby increasing the diversity of introduced base substitutions (see Figure 2a-d for the analysis of mutagenic properties).

In order to evaluate different molecular implementations of deaminase-T7RNAP fusion constructs, we performed fluctuation assays, which involve culturing cell populations for a defined time under neutral conditions, followed by subjecting the cells to stringent selective pressure which eliminates the majority of cells, leaving behind the ones that have acquired specific mutations conferring a non-lethal phenotype. This experiment enables measurement of the mutation rates in a population and, in the context of our work, provides the basis for a performance comparison of different mutagenic systems. Our new fluctuation assay utilizes a gene encoding the mitotic spindle protein She1 as a target for mutagenesis **(Figure 1b**). *She1* expressed under the control of the GAL1 promoter leads to a growth arrest phenotype when its expression is induced with galactose (2%), but does not otherwise affect growth^24^ (**Supplementary Figure 3**). This selection process eliminates the majority of cells, leaving behind only the ones that have acquired specific mutations conferring a non-lethal phenotype. The condition-dependent lethal effect enabled quantification of mutational rates by generating *she1* mutagenesis libraries in galactose-free liquid culture followed by plating on galactose-containing media to determine the fraction of cells that acquired a loss-of-function mutation. This assay was designed to provide the same functionality as the conventional yeast *can1* fluctuation assay^25^ but without the need for expensive and toxic amino acid analogues.

We first used our fluctuation assay to test the relative efficiency of mutagenesis of two T7RNAP-fused cytidine deaminases, previously evolved evoAPOBEC1^26^ and wild-type sea lamprey deaminase pmCDA1. We evaluated the performance of the designed mutator proteins in both *S. cerevisiae* EBY100 and its *ung1*-deficient mutant **(Figure 1c**). In the latter, the lack of uracil-DNA glycosylase activity prevents the excision of deaminated cytosines (uracils), impeding the activation of downstream DNA repair pathways (**Supplementary Figure 4**) and leading to the accumulation of unrepaired deoxyuridines and, ultimately, fixed C>T mutations. Both constructs showed comparable activity and approximately 100-fold improvement in *ung1*-deficient genetic background. Furthermore, we observed that increasing concentration of doxycycline inducer above 0.25 µg/ml resulted in the partial loss of mutational activity and consistently high variability between biological replicates (**Supplementary Figure 5**), even though higher doxycycline concentrations (>1 µg/ml) are typically used to achieve full induction of *tetO*-driven gene expression^27,28^ and no toxicity effects are observed below 5 µg/ml doxycycline^27^. One possible explanation for this observation is that the high rate of unrepaired DNA lesions leads to frequent polymerase stalling, which can drive an increase in the recombination rate and confer a genetic instability phenotype^29^, leading to loss of functional expression of the mutator complex. Using the same fluctuation assay, we determined that introducing G645A and Q744R point mutations into T7RNAP significantly reduced the mutation rate, even though the opposite effect had been reported for a similar system in human cells^10^ (**Supplementary Figure 6a**), and that reducing the length of the linker from 43 to 29 amino acids substantially improved the observed mutation rates (**Supplementary Figure 6b**).

Next, we designed a set of dual-mutator protein candidates that expand the attainable mutagenic spectrum. In this design iteration, we fused T7RNAP to both a cytidine deaminase (either evoAPOBEC1 or pmCDA1) and an adenine deaminase (the deaminase module of the previously reported ABE8e base editor, evoTadA^30^), forming a three-component fusion protein. We reasoned that combining both nucleoside deaminases into a single mutator protein as opposed to fusing them with separate polymerase modules, in analogy to TRIDENT^9^, would not only reduce the overall expression burden but also increase the likelihood of a mutation occurring during each event of RNA polymerase translocating alongside the DNA template (and potentially increase the total mutation rate as a single copy DNA template is likely to be saturated by the highly overexpressed polymerase). As reducing linker length was shown to be beneficial in the context of cytidine deaminase variant, the new designs were equipped with shorter linker sequences (21 and 24 amino acids).

We investigated the performance of new variants in *ung1-* and *apn1*-deficient strains. Apn1 is a major apurinic/apyrimidinic (AP) endonuclease that accounts for 97% of total AP-endonuclease activity in *S. cerevisiae*^31^. Therefore, we hypothesized that an *apn1*-deficient strain can efficiently excise uracil base from deaminated nucleosides but might show impaired fidelity when repairing the generated AP sites as the relative contribution of high-fidelity base excision repair would be diminished, while the contribution of mutagenic translesion synthesis would increase (**Supplementary Figure 4**). When the mutation rate was characterized with the *she1* assay, both dual mutator proteins performed equal or better than cytidine deaminase-based variants in all genetic contexts. The overall efficiency was approximately one order of magnitude lower in the *Δapn1* strain (**Figure 1d**). Sanger sequencing of recovered clones confirmed the predicted increase in mutational diversity relative to single mutator variants (**Supplementary Table 8**). Additionally, we tested the effect of genome integration on the performance of the evoAPO1 mutator and the evoAPO1-evoTadA dual mutator (**Supplementary Figure 7**). However, as no improvement was observed, the plasmid-based system was selected for further work in order to maximize operational flexibility. For long-term evolution experiments, the use of a genome-based system might be considered in order to reduce the risk of mutator phenotype loss due to chromosomal and episomal recombination events that have been observed for ARS/CEN plasmids^32^. Based on our initial results, we selected two strains for further characterization: EBY100 Δ*ung1*/pMut-evoAPO1_evoTadA_T7RNAP (YeastIT1), which displayed highest mutation rate, and EBY100 Δ*apn1*/pMut-evoAPO1_evoTadA_T7RNAP (YeastIT2), which was expected to provide a more diverse mutational spectrum (**Figure 1e**).

### Characterization of mutational rates and spectra in YeastIT1 and YeastIT2

Mutation rate and spectrum are the two most important factors that influence the outcome of library generation with random mutagenesis^33^. Most of the existing in vivo mutagenesis systems exhibit mutation spectra with >90% transitions and/or suffer from low mutational rates. To explore potential advantages of YeastIT1/2 strains over previously reported ones, we characterized their mutational profiles. We used a UMI-linked consensus sequencing workflow^21^, which performs an accurate inference of full-length consensus sequences, allowing characterization of not only the mutagenic rate and spectra, but also the distribution of mutational rates across the population. Initial characterization of the observed mutation distribution revealed that both strains introduce mutations throughout the gene, with mutational hotspots at the beginning and at the end of the coding sequence. The observed mutations included all four mutations that correspond to cytidine and adenosine deamination (A>G, T>C, G>A, C>T), indicating activity of the two deaminase domains on both coding and template strand (**Figure 2a**). YeastIT1 exhibited substantially higher mutation rates, averaging 4.19 point mutations/gene (1.38 mutations/kb/day), while YeastIT2 had lower rates of 0.26 point mutations/gene (0.09 mutations/kb/day) (**Figure 2b**). Notably, mutational frequencies observed for the YeastIT1 system do not follow a negative binomial distribution that would be expected for a culture with homogenous mutator phenotype, but rather display a bimodal distribution. This observation can be attributed to the steady-state bimodal response of the reverse tetracycline transactivator^34^ that controls the expression of the mutator protein arising from its cooperative binding to multiple *tetO* binding sites^35^. The mutation rate in the YeastIT2 library was too low to provide insights about the bimodal pattern of mutator protein expression.

**Figure 2.**
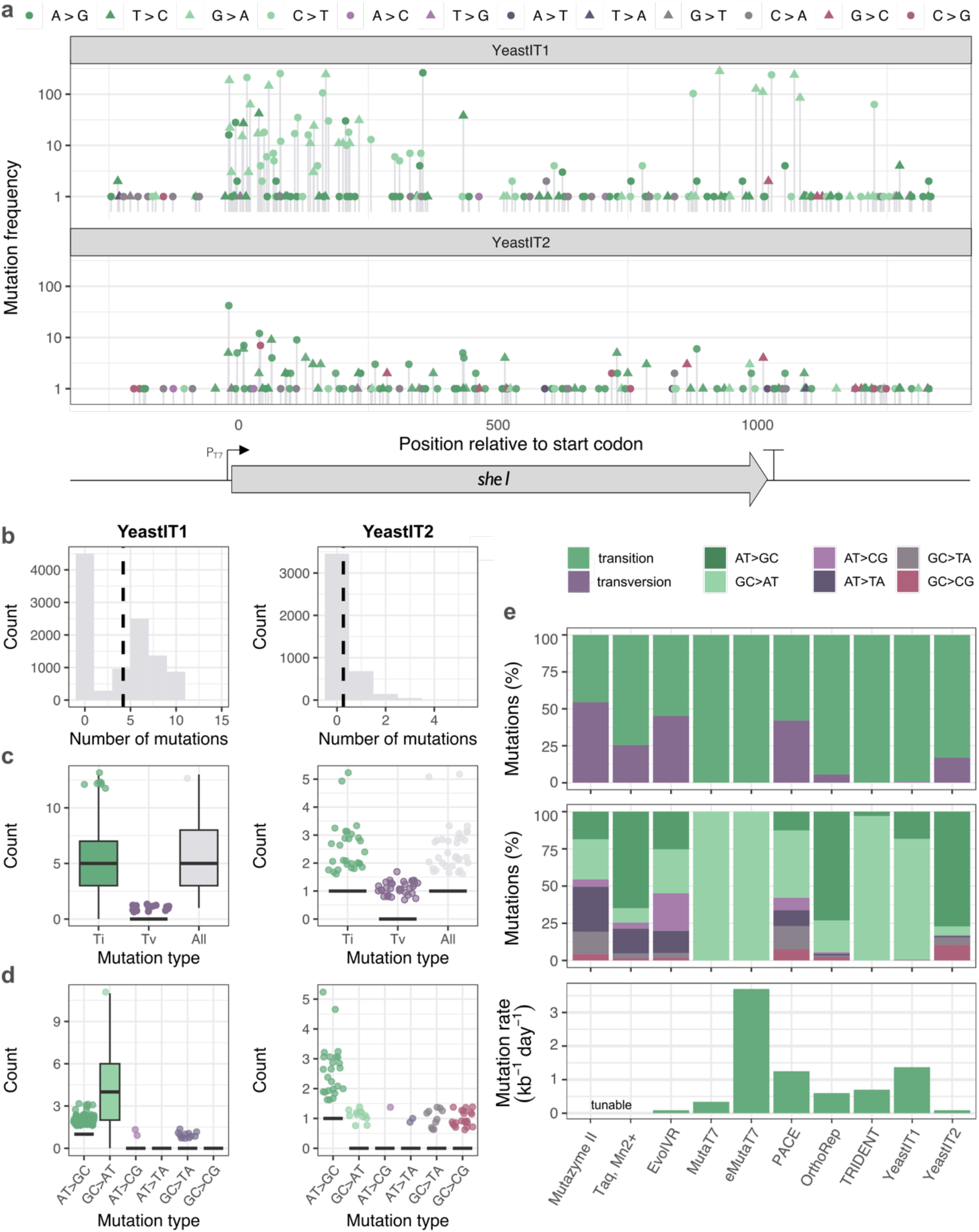
Characterization of mutational frequencies and spectra in variant populations derived from YeastIT1- and YeastIT2-generated *she1* libraries. (**a**) Distribution of mutation types at the *she1* target locus. Position 0 indicates the start codon. Six mutation types are indicated with colour codes. Circles correspond to mutations with deaminated base on the coding strand. Triangles correspond to mutations with deaminated base on the template strand. (**b**) Distribution of mutation counts in YeastIT1-derived (*left*) and YeastIT2-derived (*right*) libraries. Mean mutation count for each sample is indicated with a dashed line. (**c**) Distribution of transitions (Ti) and transversions (Tv) in YeastIT1 (*left*) and YeastIT2 (*right*) variants. YeastIT2 generates mutations at a lower rate but achieves a higher number of transversions. (**d**) Distribution of 6 mutation types in YeastIT1 (*left*) and YeastIT2 (*right*) variants. Histograms (**b**) show distribution of mutational frequencies in all sequences (10,503 for YeastIT1 and 4,336 for YeastIT2), while boxplots (**c**,**d**) show distribution of mutation types in unique variants (473 unique variants with 2,505 mutations for YeastIT1, 148 unique variants with 201 mutations for YeastIT2). In **b**, **c** and **d** only mutations occurring within *she1* open reading frame were considered. In boxplots, the centre line indicates the median, the box is the interquartile range (IQR) and whiskers indicate the most extreme value within 1.5 IQR. (**e**) Comparative analysis of mutational frequencies and spectra in different in vivo and in vitro mutagenesis methods.

Next, we set out to establish whether the lower mutation rates of YeastIT2 have a substantial negative effect on the library diversity. Accurate quantification of diversity in libraries generated through in vivo mutagenesis is challenging due to confounding factors such as mutational hotspots, complex mutational frequency distributions, and presence of lineages with phenotypically diverse mutator behaviours. To estimate library size based on the UMIC-seq data, we first looked at the relationship between mutational count and sequence cluster count for each unique variant in the library to exclude the possibility of non-existing variants present in the data due to sequencing errors (**Supplementary Figure 8a**). We observed no correlation between these two parameters, indicating that the identified mutation events are truly present in the population. This analysis also confirmed that both YeastIT1 and YeastIT2 libraries were heavily undersampled as most variants can be assigned to a single UMI cluster. We then estimated the upper-bound limit of library size for each method by assuming that every position is mutated with equal likelihood, every amino acid substitution is equally likely, and no two mutation events occur in a single codon (while those assumptions do not precisely reflect the reality, they have conventionally be used for estimating diversity in error-prone PCR libraries^36–38^). Probabilistic analysis resulted in estimates of 7.8×10^5^ and 1.8×10^4^ unique variants per million cells (corresponding to approximately 0.1 mL of cell culture at OD of 1) for YeastIT1- and YeastIT2-specific mutation rates, respectively **(Supplementary Table 9**). Both theoretical libraries covered approximately 1/3 of mutants with a single amino acid substitution (achieving full coverage of variants with a single non-synonymous mutation), with YeastIT1 covering 1.1% of sequence space of mutants with two non-synonymous mutations, and YeastIT2 covering 0.07% (**Supplementary Figure 8b**). This difference in sequence space coverage for variants with two non-synonymous mutations can be mitigated by scaling-up the screening efforts. This suggests that, when ultra-high-throughput screening methods such as flow cytometry or droplet microfluidics are available, an unbiased mutational spectrum becomes the key feature to prioritize for a successful directed evolution campaign.

Further analysis of mutation counts for each type of nucleotide substitution revealed that YeastIT2 mutagenesis leads to a more diverse spectrum of mutations than YeastIT1, with the significant increase in transversion mutations from 0.6% to 16.9% (**Figure 2cd**). This increase in transversions occurs exclusively at G/C positions, which is to be expected as Ung1 has been reported to be highly specific for uracil^39^ and no promiscuous activity on hypoxanthine has been reported. We further expanded our analysis to compare the performance parameters of YeastIT1 and YeastIT2 with those previously reported for other in vivo mutagenesis methods (see **Supplementary Table 7** for data sources) and found that YeastIT2 exhibits a higher proportion of transversions than alternatives such as MutaT7, eMutaT7, OrthoRep, and TRIDENT (**Figure 2e**). PACE and EvolVR, while still favouring transitions, showed less biased mutagenic spectrum than YeastIT strains.

Differences in obtainable mutagenesis spectra between methods translate into prominent differences in amino acid substitution probabilities and protein sequence space accessibility, therefore making the choice of genetic diversification method a key determinant in the success of the directed evolution campaign. Therefore, we explored how the mutagenesis spectra observed in DNA libraries generated with YeastIT strains, previously developed in vivo mutagenesis methods (MutaT7, eMutaT7, TRIDENT, PACE, OrthoRep), and more widely adapted in vitro mutagenesis methods (error-prone PCR with Taq polymerase and Mn^2+^ supplementation or with Mutazyme II kit) affect the amino acid diversity.

To establish a benchmark for all methods, we first sought to determine the optimal mutagenic spectrum for effective genotype diversification. Living organisms display non-random mutational patterns, with transitions almost universally occurring at higher rates than transversions (usually 2-to 5-fold increase)^40–43^. The exact underlying reason for this phenomenon remains unclear with one hypothesis suggesting that transition bias may be partially attributed to transitions being more likely to lead to conservative rather than radical amino acid substitutions^44^, with some studies providing contradictory evidence^45^, and others demonstrating that the deleterious impact of transversions is highly species-dependent as evidenced by large variations in amino acid ‘exchangeability’ (fixation probability of a given amino acid substitution) among species^46^. The evolutionary benefit of minimizing phenotypic effects of mutations is further supported by the preference in codon assignment for both synonymous mutations and substitutions to amino acids with similar chemical properties, particularly polarity^47,48^. While, in a directed evolution context, a system with higher likelihood of perturbative mutations than observed in Nature could be advantageous, achieving this would require modifying the canonical genetic code. Therefore, we considered a theoretical system with equal probability of each DNA mutation as a practically feasible approximation of an optimal scenario for benchmarking purposes.

Analysis of the relative amino acid substitution probabilities revealed that all methods perform worse than the theoretical benchmark with only PACE and in vitro mutagenesis methods surpassing YeastIT2 in terms of the total number of probable substitutions and uniformity of the probability distribution (**Figure 3ab, Supplementary Figure 9**). Given the probability threshold of 0.05, YeastIT2 is likely to perform 80 out of the total of 153 amino acid substitutions possible with a single nucleotide exchange and YeastIT1 is likely to perform 57 out of 153 substitutions, while PACE can perform 150 substitutions (albeit with a more skewed probability distribution than the ideal system, leaving room for further improvement).

To further determine possible differences in each method’s preference for transitioning between the chemical properties brought about by various amino acids, we constructed simplified matrices (**Figure 3c**) in which amino acids are categorised by their molecular recognition properties: polar, non-polar, basic, and acidic. This framework allows quantification of the number of single nucleotide exchanges that enable each shift in chemical space. Firstly, we reasoned that it is important to balance the exploration of sequence space with the risk of introducing deleterious mutations. Since most functional residues in proteins are polar or charged (the seven amino acids most frequently involved in catalytic roles are C, H, D, K, S, T and Y)^49^, we evaluated the potential for functional innovation by quantifying the ratio of substitutions into polar or charged residues relative to substitutions into nonpolar residues (**Figure 3d**). We found that this ratio was particularly high for YeastIT2 and surpassed the methods with more diverse mutational spectra (e.g., 1.44-fold higher than for Mutazyme II, 1.53-fold higher than PACE). This indicates that YeastIT2 preferentially introduces more new residues with functional potential while making relatively few substitutions to nonpolar residues, which are more likely to disrupt function. Secondly, we observed that YeastIT2 had the highest ratio of substitutions resulting in negative carboxylates (D, E) compared to substitutions resulting in amino acids with positive ammonium groups (H, K, R) among all methods, while YeastIT1 ranks the highest for the reciprocal conversion (**Figure 3d**). This behaviour could be exploited for surface charge engineering of proteins to alter their pH profile, salt tolerance, or stability. For example, the pH optimum of a xylanase was shifted towards the alkaline region with concomitant improvement of thermostability by introducing 5 surface arginines^50^; similarly, a carbonic anhydrase was engineered for extreme halotolerance by introducing 18 acidic residues on its surface^51^.

**Figure 3.**
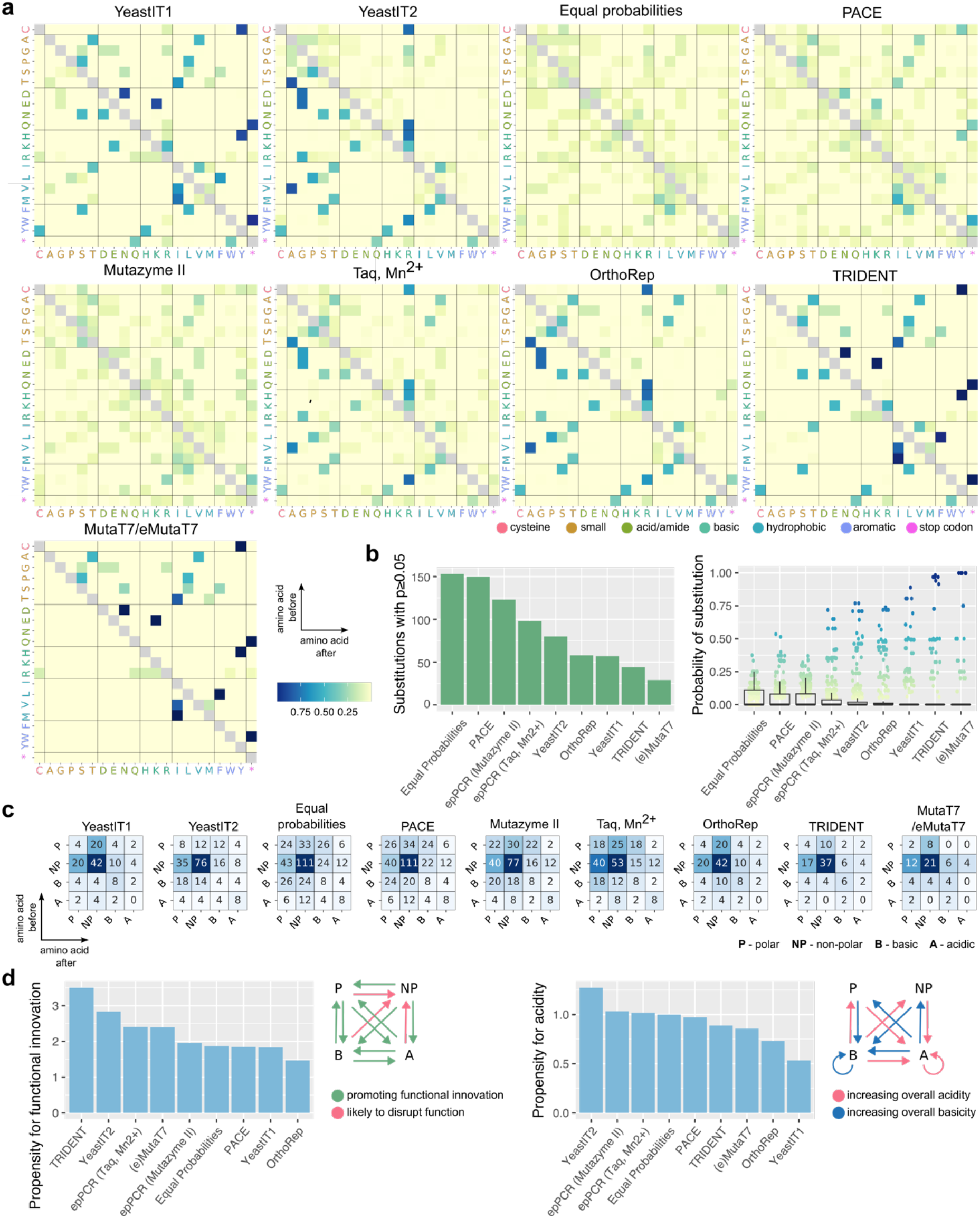
Amino acid substitution probabilities and molecular recognition feature accessibility of random mutagenesis methods. (**a**) Experimental amino acid substitution matrices of random mutagenesis methods and an idealized system performing every mutation with equal probability. Matrices display relative probability of each substitution for a given amino acid assuming even distribution of all possible codons (more detailed codon matrices are shown in **Supplementary Figure 9**). EvolVR was excluded in this comparison as the method does not allow full-length gene mutagenesis. Synonymous mutations were excluded from visualisation. (**b**) Overview of amino acid substitution probability statistics for mutagenesis methods summarizing the total number of substitutions with p≥0.05 (*left*) and distribution of probabilities for all substitutions (*right*, individual values for substitutions with p≥0.05 shown as dots). (**c**) Matrices describing access to molecular recognition features, categorised as P (polar: S, T, Y, N, Q, C), NP (non-polar: A, V, I, L, M, F, W, G, P), B (bases: K, L, H), A (acids: E, D), summarizing the ‘chemical space’ explorable by different random mutagenesis methods and an idealized system performing every mutation with equal probability. The total number of codon substitutions with p≥0.05 for each transition between chemical spaces is given on the heatmap. (**d**) Overview of changes in molecular recognition, highlighting trends for mutagenesis methods including the ratio of mutation types promoting functional innovation to mutations likely to disrupt function (*left*) and the ratio of acid (carboxylate: E, D)-increasing to base (K, L, H)-increasing mutations (*right*). Diagrams next to the graphs indicate the substitution types used for calculating each ratio.

To explore mutation trajectories in long-term continuous evolution experiments, where multiple mutations can occur at each position over time, we further analysed the mutational spectra to generate force-directed graphs which visualize relationships between codons and, thus, amino acid diversity accessible with each method. Force-directed graphs (**Figure 4a)** illustrate relationships between 64 codons, which are represented as nodes and connected by edges whose opacity indicates the probability of substitution. This visualisation allows a classification into two categories: the first class (YeastIT2, PACE, error-prone PCR) features highly interconnected network while the second class (OrthoRep, MutaT7, eMutaT7 YeastIT1, and TRIDENT) displays the formation of 8 clusters with high probabilities of substitutions within the cluster and low or no likelihood of escape. The lack of interconnectedness observed in the second class indicates limited access to diversity (at certain positions, substitutions will only occur within a narrow set of few codons regardless of the length of the experiment), and, thus, a constrained capacity for long-term sequence space exploration. Among those methods, only OrthoRep features bridge nodes which enable rare but crucial events of traversing between clusters, furthering the attainable sequence diversity. On the other hand, YeastIT2, PACE, or error-prone PCR have no propensity for clustering, indicating that they can generate more diverse libraries and enable long adaptive walks with lower susceptibility to be trapped in local fitness maxima.

**Figure 4.**
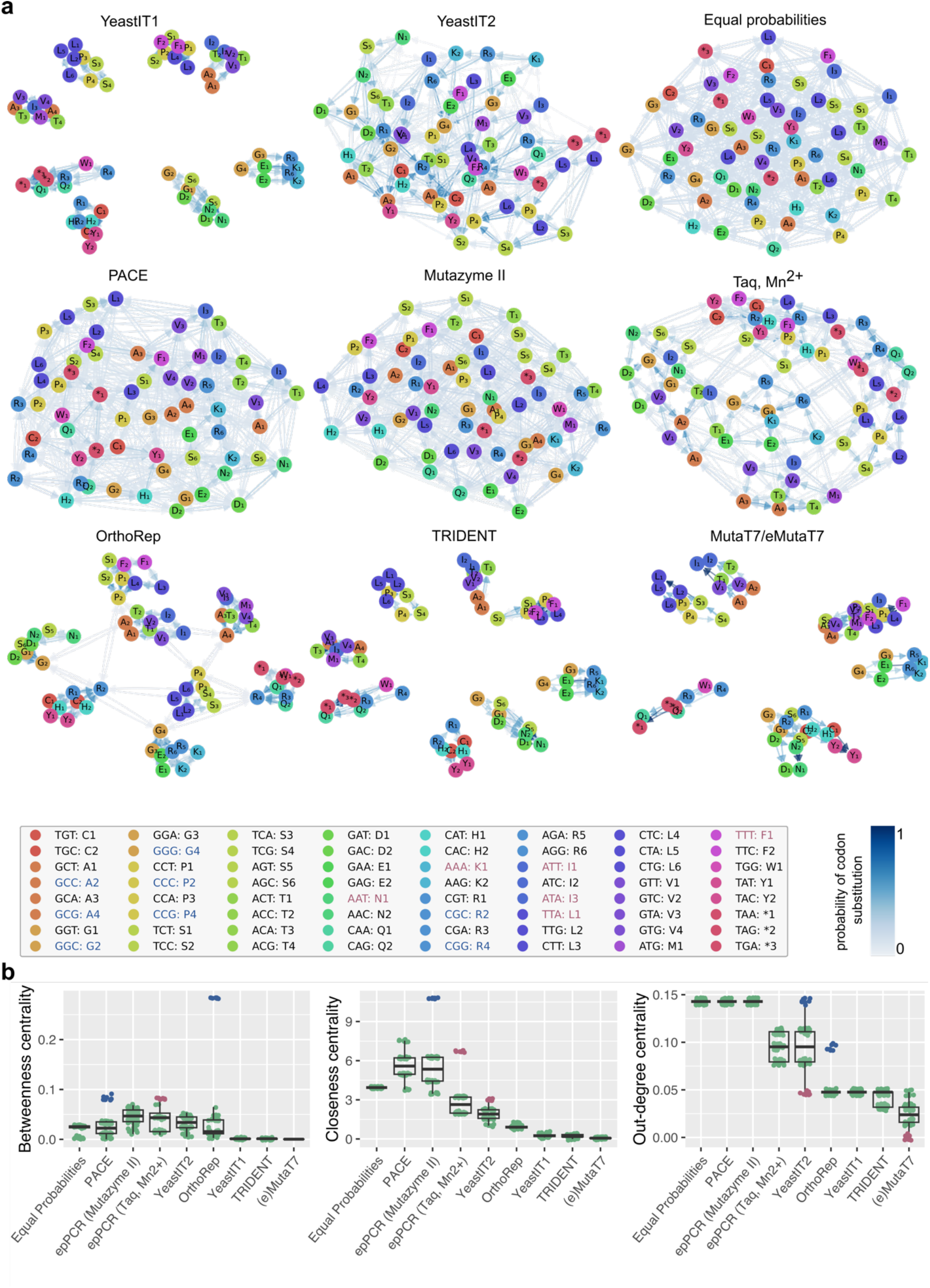
Codon substitution probabilities of random mutagenesis methods visualized as force-directed graphs. (**a**) Force-directed graphs representing the diversity of experimentally observed random mutagenesis libraries generated with various methods and an idealised system performing every mutation with equal probability. Graphs were constructed using a Fruchterman-Reingold algorithm and display frequently interconverting codons as clusters. Nodes represent codons, edges represent substitutions (one-base interconversions), edge opacity indicates the likelihood of the substitution relative to all other substitutions for a given codon. For clarity, edges corresponding p<0.025 are not displayed. The box contains shorthand codes for triplet codons and their corresponding amino acids that are annotated in the force-directed graphs. (**b)** Overview of the force-directed graph statistics – node centrality parameters describe connectivity and importance of individual amino acids in the network. For any method with an unbalanced mutation spectrum, these parameters can provide a quantitative measure of a codon’s suitability as a starting point for sequence space exploration. Betweenness centrality refers to a sum of all-pair edge-weight-normalized shortest paths that pass through a node and describes its importance for the flow in the network, closeness centrality refers to a reciprocal of the edge-weight-normalized shortest path to all reachable nodes scaled by the fraction of reachable nodes and depends on the relative mutational frequency of a codon rather than the diversity of its mutational spectrum, and out-degree centrality refers to the fraction of all possible outgoing connections and prioritizes codons with most balanced mutational spectrum. The outlier codon groups, characterized by exceptionally high or low mutation frequencies or mutational diversity, are highlighted and mapped to the legend box in **a**. High-scoring outliers may be preferentially considered in the initial gene design. Additionally, codons with low out-degree centrality score are more likely to be trapped in local fitness maxima when chosen in the design or encountered during mutagenesis.

Additionally, we wanted to establish whether, given the mutational spectrum of each method, encountering certain codons might be particularly beneficial or disadvantageous during the directed evolution campaign that employs it. To determine their enabling role in evolution, node centrality parameters - betweenness centrality (i.e. importance for the flow in the network), closeness centrality (shortness of paths to other nodes), and out-degree centrality (number of outgoing connections) - were determined for each force-directed graph (**Figure 4b**) and the codon identity of outlier nodes was identified. High-scoring codons were observed for all three parameters and low-scoring codons were observed for out-degree centrality. While the graph parameters do not directly correlate with specific mutational parameters, closeness centrality tends to be more influenced by relative mutational frequency of each codon, whereas out-degree centrality is more influenced by mutational spectrum attainable for each codon. This can be observed most clearly in the YeastIT2 dataset, where AT-rich codons constitute a high-scoring outlier group for the former parameter because they are mutated more frequently and a low-scoring outlier group for the latter because the occurring mutations are less diverse. These considerations describe the ability of the mutational spectrum introduced to lead to different potential functional solutions during cycles of library synthesis and screening. We conjecture that these properties are useful to consider during the design of initial gene to be evolved, where high-scoring codons are likely to be preferred, and when assessing if a particular mutation identified in the improved variant is likely to be located at local fitness maximum (low-scoring codons are more prone to this behaviour).

### Evolution of GFP-binding DARPin 3G146

*S. cerevisiae* has been employed as a host in a diverse range of directed evolution campaigns (metabolic engineering^52,53^, enzyme engineering^54–56^, binder engineering^57–59^) with one of its most important applications being affinity maturation of therapeutically relevant antibodies and other binding scaffolds^60,61^, a laboratory equivalent of the somatic hypermutation process of the adaptive immune system. To showcase the applicability of YeastIT for binder affinity maturation, we engineered yeast to display the previously identified GFP-binding DARPin 3G146 (reported affinity of 156 nM^17^) and performed 2 rounds of iterative YeastIT library generation and FACS-based screening (**Figure 5a, Supplementary Figure 2**), successfully identifying several clones that showed improved affinity in secondary flow cytometry assays (**Supplementary Figure 10, Supplementary Figure 11**). We selected 13 clones for sequencing and identified 7 unique variants, with 2 mutations (G150S and G237S) within the DARPin-coding region and other mutations occurring in N-terminally fused Aga2 domain, signal peptide, linker region containing HA tag, and C-terminally fused c-myc tag (**Supplementary Figure 12**). Out of those, we selected variant 2-G4 (G150S mutation) and variant 2-H3 (G237S mutation) for quantification of apparent binding affinity using on-yeast surface titrations (**Figure 5b**, **Supplementary Figure 13**). Furthermore, to determine whether the improvements in binding affinity can be attributed to a yeast display-specific effect, we expressed the variants in a soluble form without the Aga2 domain (numbering shifted to G54S in 2-G4 and G141S in 2-H3 due to removal of N-terminus) and performed further characterization with ELISA and BLI (**Figure 5b**, **Supplementary Figure 14, Supplementary Figure 15**). Affinity improvement was confirmed by both in vitro measurements in 2-H3 (15-fold *K_d_* improvement in BLI), while 2-G4 showed no detectable improvement in ELISA and only minor improvement in BLI. As the variant 2-G4 has a mutation within the N-terminal capping repeat located in close proximity to the Aga2 domain, we speculate that this residue might be involved in the interactions with Aga2 and affect the spatial orientation of the yeast-displayed protein. Consequently, it results in a much higher affinity improvement in the context of being displayed on yeast than in a solution.

**Figure 5.**
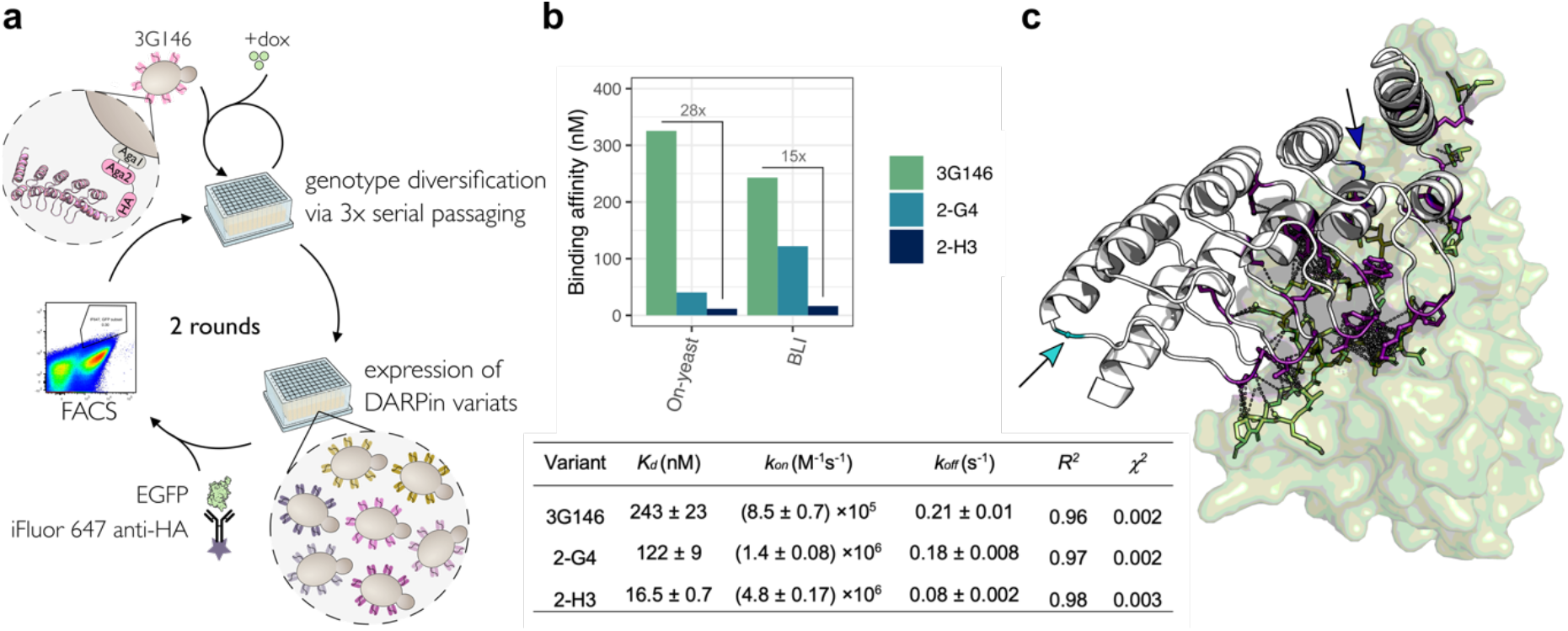
Directed evolution of GFP-binding DARPin 3G146 for improved binding affinity. (**a**) Overview of experimental approach for in vivo library generation and FACS-based screening. Two rounds of genetic diversification and screening with decreasing EGFP concentration were performed. (**b**) Binding affinity characterization using on-yeast titration assay (reflecting the screening conditions), showing 28-fold improvement for the variant 2-H3 and 8-fold improvement for the variant 2-G4, and using BLI, showing 15-fold improvement for the variant 2-H3 and 2-fold improvement for the variant 2-G4 (*top*). Summary table of BLI-derived *K_d_*, *k_on_*, and *k_off_* values (*bottom*, values are reported as mean (±standard error) of 3 measurements at different DARPin concentrations). BLI traces are shown in **Supplementary Figure 15**. (**c**) AlphaFold2-predicted structure of 3G146:EGFP complex with a G141S position mutated in 2H-3 indicated in blue and a G54S position in 2-G4 indicated in cyan. Both residues are not located directly at the binding interface. Residues located within 4.5 Å of the binding partner are shown as balls and sticks with their potential interactions shown as dotted lines.

Engineering high-affinity binders is often accomplished through rational approaches where residues located at the binding interface are selectively targeted for mutagenesis^62,63^. Given the experiment-defying vastness of sequence space, this strategy can be rationalized as increasing the chance to discover mutations that elicit a (positive) functional effect. In the context of DARPin engineering, this is typically done by employing a consensus design approach, wherein positions exhibiting variability in natural ankyrin repeat proteins are randomized, while residues conserved in Nature are left unchanged^17,64–66^. Interestingly, the affinity-improving mutations identified in this study were located outside the variable regions (**Figure 5c**, **Supplementary Figure 16**): 2-H3 displayed a mutation at position G16 of the third ankyrin repeat, a residue highly conserved in Nature (60-80%)^64^, while 2-G4 had a mutation at position G25 in the N-terminal capping repeat. This finding highlights the potential of random mutagenesis to yield variants with improvements conferred by mutations occurring away from the typically targeted sites, thereby providing access to more diverse sequence repertoire. Similar observations were previously reported for antibody fragments in a systematic study that compared the performance of error-prone PCR and rational CDR-targeted libraries in affinity maturation, demonstrating similar improvements in *K_d_* for best variants obtained with both methods, with error-prone PCR yielding variants with beneficial mutations located outside of the CDR regions^67^.

To investigate whether prediction of the binding affinities in protein-protein interactions would be possible as an alternative to experimental evolution, we applied several available prediction tools (TopNetTree^68^, SAAMBE-3D^69^, mCSM-PPI2^70^, FoldX^71^, SSIPe^72^), to determine if the experimentally discovered variant could be predicted in silico. However, none of the tools were able to accurately predict the direction or magnitude of the binding affinity change caused by the point mutation in the 2H-3 variant (**Supplementary Figure 17**). This highlights the limitations of currently available methods in predicting the effects of single point mutations on binding (as recently pointed out by others^73,74^) and emphasizes the need for improved high-throughput methods in experimental protein engineering to generate large datasets that can fuel the development of next-generation prediction algorithms.

## Discussion

In vivo mutagenesis systems hold the promise of making protein evolution automated, scalable and user-friendly. Their impact is crucially affected by factors such as ease of implementation, accessibility to a wide mutational spectrum and compatibility with high-quality and high-throughput assays. Our newly developed mutator strain addresses these requirements effectively: (i) YeastIT uses a widely adopted yeast host known for its protein display and secretory capacity as well as straightforward genetic manipulation; (ii) it offers a more diversified mutagenesis spectrum compared to most existing in vivo alternatives (the methods that report more balanced mutagenesis spectra suffer from major limitations – PACE workflow requires coupling the phenotype of interest to selection pressure and a relatively complex experimental set-up, while EvolVR targets only a short nucleotide window instead of full-length gene and therefore cannot be considered a truly continuous evolution system); and (iii) it can be effectively integrated with ultra-high-throughput assays such as flow cytometry (as in this work), droplet microfluidics (where quantitative picoliter assays for all seven Enzyme Commission classes are available^75^), or growth selection. Combining yeast expression with those assays has already established itself as a powerful tool for optimizing binding affinity in antibody fragments^76^, antibody mimics^59^ or T-cell receptors^77^ and activity of enzymes such as β-glucosidase^78^, caffeine demethylase^79^, horseradish peroxidase^80^, or glucose oxidase^81^.

The successful demonstration of our directed evolution campaign using YeastIT resulted in the identification of a mutant with a 15-fold improvement in affinity through a single mutation. This outcome suggests that the individual components of YeastIT are well integrated, validating its utility. However, applying in vivo mutagenesis to longer evolution campaigns will be even more attractive due to the cumulative benefit of eliminating re-cloning steps between rounds. In such long evolution campaigns where the operational requirements for successful applications demand many mutations (such as, for example, the evolution of a transaminase to catalyse the amination of prositagliptin ketone for the synthesis of sitagliptin that required 11 rounds of directed evolution and resulted in a total of 27 mutations^82^), the benefits of an in vivo platform with high throughput and wide mutational spectrum, which provide access to large and diverse libraries, become even more evident.

A major advantage of in vivo mutagenesis is its ability to expand the size of screened libraries. In contrast to the limitations faced by in vitro-generated libraries expressed in yeast, which typically feature a diversity of 10^7^-10^8^ variants due to transformation efficiency limitations (up to 10^10^ reported with extensive scaling-up efforts^83^), our platform enables the generation of the diversity that matches the screening throughput of a FACS assay preceded by magnetic bead enrichment (10^10^ for a routine experiment with further scale-up possible^84^). This places yeast libraries on par with phage libraries in terms of size, while also providing access to unique advantages offered by yeast surface display such as the facilitation of post-translational modifications^85^, possibility of quantitative analysis of dissociation kinetics^85^, and superior diversity of the isolated immune repertoire from identical libraries (implying higher fraction of the library being functional)^86^.

Our YeastIT2 strain exemplifies the principle of enhancing mutational diversity by integrating a nucleoside deaminase mutator^7^ with the host’s error-prone DNA repair machinery. This integration resulted in a mutator strain that achieved a 28-fold improvement in transversions (16.9% vs 0.6%) relative to the only previous implementation of this strategy^9^. We anticipate that with further engineering of the DNA repair system, even more balanced mutagenic spectra and improved mutation rates will be achieved in the future. For instance, one potential approach could involve an overexpression of a DNA glycosylase capable of effectively excising hypoxanthine, thereby enabling the conversion of deaminated adenines into pyrimidines and leading to enhanced mutational diversity. Another approach to increase the mutation rate and distribute mutations more evenly across the gene could involve incorporating T7 promoters throughout the gene in designed intronic regions, which would later be excised.

The comparative analysis of evolutionary trajectories achievable by YeastIT and other mutagenesis methods presented in **Figure 3** and **Figure 4** reveals substantial differences in their sequence space accessibility and evolutionary potential as well as highlights the significance of choosing the starting point in directed evolution. While previous studies have already demonstrated that both the mutagenesis method^67^ and the codon usage during the input gene design^87^ influence the evolutionary outcomes, our systematic comparison of all methods for which necessary data were available serves as a valuable tool for making informed decisions in experimental design during directed evolution efforts. Additionally, our analysis framework will prove useful in benchmarking future mutagenesis systems to YeastIT and other methods.

Finally, tracking evolutionary trajectories through next-generation sequencing can provide further insights into protein evolution. The lowered technical barrier due to working with yeast as a robust host and obviating the need for cloning in mutagenesis, coupled with the freedom to define directionality and adaptive versus neutral (non-adaptive) evolution regimes based on carefully controlled selection pressure, will lead to much more rapid access to multi-round deep sequencing datasets, which can be interpreted by emerging AI/ML methods^88^.

## Reporting summary

Further information on research design is available in the Nature Portfolio Reporting Summary linked to this article.

## Code availability

All code is available at https://github.com/fhlab/YeastIT.

## Data availability

Raw sequencing data are deposited in the European Nucleotide Archive under the accession code PRJEB65953.

## Acknowledgements

This work was supported by the European Union’s Horizon 2020 research and innovation programme via the Marie Curie training network EVOdrops (to MN, 813786). FH is an ERC Advanced Investigator (695669). We thank Paul Dupree and members of the Hollfelder group for helpful discussions.

## Author Contributions

MN, KF, PK and FH designed research. MN and KF performed experiments. MN and MP performed bioinformatic analyses. MN and FH wrote the paper with input from all authors.

## Competing Interests

None

## Notes

### Competing Interest Statement

The authors have declared no competing interest.

