## Supplementary Information for "YeastIT: Reducing mutational bias for in vivo directed evolution using a novel yeast mutator strain based on dual adenine-/cytosine-targeting and error-prone DNA repair"

**Supplementary Table 1. Strains used in this study**

| Name | Genotype/Features | Reference |
|---|---|---|
| *E. coli* BL21(DE3) | *fhuA2 [lon] ompT gal (λDE3) [dcm] ΔhsdS* | New England Biolabs |
| *E. coli* NEB5-alpha | *fhuA2Δ(argF-lacZ)U169 phoA glnV44 Φ80Δ(lacZ)M15 gyrA96 recA1 relA1* | New England Biolabs |
| *S. cerevisiae* EBY100 | *MATa AGA1::GAL1-AGA1::URA3 ura3-52 trp1 leu2-Δ1 his3-Δ200 pep4::HIS3 prbΔ1.6R can1 GAL* | ATCC MYA-4941 |
| *S. cerevisiae* EBY100 *Δung1* | *S. cerevisiae* EBY100 *ung1Δ::KANMX* | This work |
| *S. cerevisiae* EBY100 *Δapn1* | *S. cerevisiae* EBY100 *apn1Δ::KANMX* | This work |
| *S. cerevisiae* EBY100 *Δung1 HO::EVOAPO1* | *S. cerevisiae* EBY100 *ung1Δ::KanMX HO::evoAPO1_T7RNApol-LEU* | This work |
| *S. cerevisiae* EBY100 *Δapn1 HO::YEASTIT2* | *S. cerevisiae* EBY100 *apn1Δ::KanMX HO::evoAPO1_evoTadA_T7RNApol-LEU* | This work |

### Supplementary Table 2. Plasmids used in this study

| Name | Features | Reference |
|---|---|---|
| HO-poly-KanMX4-HO | pBR322 origin of replication, ampicillin resistance, integration of KanMX cassette into HO locus in *S. cerevisiae* | [1] |
| pMut-evoAPO1_T7RNAP | ARS4/CEN6 yeast origin of replication, Leu auxotrophic marker, fusion of evoAPOBEC1 and T7RNAP expressed under the control of $tetO_6$ operator sequence, expression of rtTA(G72P) transactivator under the control of TDH3 promoter, ColE1 origin of replication, ampicillin resistance | This work |
| pMut-evoAPO1_evoTadA_T7RNAP | Derived from pMut-evoAPO1_T7RNAP, evoTadA module inserted | This work |
| HO-evoAPO1_T7RNAP-LEU-HO | Derived from HO-poly-KanMX4-HO-and pMut-evoAPO1_T7RNAP, integration of rtTA-evoAPO1_T7RNAP-LEU cassette into HO locus | This work |
| HO-evoAPO1_evoTadA_T7RNAP-LEU-HO | Derived from HO-poly-KanMX4-HO and pMut-evoAPO1_evoTadA_T7RNAP, integration of rtTA-evoAPO1_evoTadA_T7RNAP-LEU cassette into HO locus | This work |
| pASK-IBA63b+ | ColE1 origin of replication, ampicillin resistance, preparation of *she1* library for UMIC-seq | IBA GmbH |
| pGoi-Aga2-3G146 | ARS4/CEN6 yeast origin of replication, Trp auxotrophic marker, galactose inducible expression of 3G146 DARPin with N-terminal Aga2 domain and HA-tag, and C-terminal c-myc tag, contains $P_{T7}$ sequence integrated downstream of galactose-inducible promoter, ColE1 origin of replication, kanamycin resistance | This work |
| pMut-pmCDA1_T7RNAP | ARS4/CEN6 yeast origin of replication, Leu auxotrophic marker, fusion of pmCDA1 and T7RNAP expressed under the control of $tetO_6$ transactivator, ColE1 origin of replication, ampicillin resistance | This work |
| pMut-pmCDA1_evoTadA_T7RNAP | Derived from pMut-pmCDA1_T7RNAP, evoTadA inserted as a middle domain | This work |
| pMut-pmCDA1_T7RNAP* | Derived from pMut-pmCDA1_T7RNAP, 2 point mutations in T7RNAP module | This work |
| pMut-pmCDA1_29aa_T7RNAP | Derived from pMut-pmCDA1_T7RNAP, linker sequence altered | This work |
| pMut-pmCDA1_51aa_T7RNAP | Derived from pMut-pmCDA1_T7RNAP, linker sequence altered | This work |
| pMut-pmCDA1_P2A_T7RNAP | Derived from pMut-pmCDA1_T7RNAP, P2A peptide integrated between modules | This work |
| pGoi-she1 | Derived from pGoi-Aga2-3G146. Aga2-3G146 insert was replaced with she1 | This work |
| pGoi-she1Δ | Derived from pGoi-she1, she1 sequence was replaced with C-terminally truncated she1ΔH246-R337 variant | This work |
| pGoi-Aga2-3G146-2G4 | Isolated after 2 rounds of directed evolution of pGoi-Aga2-3G146 | This work |
| pGoi-Aga2-3G146-2H3 | Isolated after 2 rounds of directed evolution of pGoi-Aga2-3G146 | This work |
| pET-28a(+) | pBR322 origin of replication, kanamycin resistance, expression of gene insert under control of $P_{T7}$-lac promoter, expression of LacI under control of the $P_{lacI}$ promoter | Merck |
| pET-28a-3G146 | Derived from pET-28a(+), expression of DARPin 3G146 with N-terminal HA-tag and C-terminal $His_6$-tag | This work |
| pET-28a-3G146-2G4 | Derived from pET-28a(+),expression of DARPin 2-G4 with N-terminal HA-tag and C-terminal $His_6$-tag | This work |
| pET-28a-3G146-2H3 | Derived from pET-28a(+), expression of DARPin 2-H3 with N-terminal HA-tag and C-terminal $His_6$-tag | This work |

### Supplementary Table 3. Sequences of primers used in this study

| Primer | Sequence |
|---|---|
| F1 | ATGCTTTTAAAGAGTTTGGAAATCGAGACCTGCATATGCAATAGTAATATGACATGGAGGCCCAGAATAC |
| R1 | ATCCATAATGATATATATTATCAGAAGCTGTACACAAGCCGTTCACATACCAGTATAGCGACCAGCATTC |
| F2 | TTAGTTACGTACGTTGAGATAATCTACAAAAATTGATTACGTATTTAAAATTCTTCTCGCTTCTCAGACATG<br>GAGGCCCAGAATAC |
| R2 | GACACCTAGCTTTGTTAGATCTGCTGTCTCGAAATACAAATTTGGTGCGCACATGTCAGGTGCCCAGTAT<br>AGCGACCAGCATTC |
| F3/F4 | TACATCGAACTGGATCTCAACAGCGNNNYRNNNYRNNNYRNNNYRNNNYRGATAC<br>NNNYRNNNYRNNNYRNNNYRNNNYRTTCTGGGGTAATTAATCAGCGA |
| R3 | CTTACTTCTGACAACGATCGGAGGACTCGATTCCGTTTGTAGTCGTCTGTGTGTATAGAAGTGAATAGCTATATAAAG |
| R4 | CTTACTTCTGACAACGATCGGAGGACGAGTCTTGTGTCCCAGTTACCAGGGTGTATAGAAGTGAATAGCTATATAAAG |
| F5 | TAACTTTAAGAAGGAGATATACCATGCCATGGTACATCGAACTGGATCTCAACAG |
| R5 | CGCAAGCTTGTCGACGGAGCTGAGCTCCTTACTTCTGACAACGATCGGAG |

**Supplementary Table 4. Mutator protein sequences used in this study**

| Name | Sequence[a] |
|---|---|
| evoAPO1-T7RNAP | MKRTADGSEFESPKKKRKVSSKTGPVAVDPTLRRRIEPHEFEVFFDPRELRKETCLLYEINWGGRHSIWRHTSQNTNKHV<br>EVNFIEKFTTERYFCPNTRCSITWFLSWSPCGECSRAITEFLSRYPNVTLFIYIARLYHLANPRNRQGLRDLISSGVTIQ<br>IMTEQESGYCWHNFVNYSPSNESHWPRYPHLWVRLYVLELYCIILGLPPCLNILRRKQSQLTSFTIALQSCHYQRLPPHI<br>LWATGLKSGGSSGGSSGGGSETPGTSESATPESGGSDYKDDDDKGSLIKGMNTINIAKNEFLEPPKKKRKVEFSDIELAA<br>IPFNTLADHYGERLAREQLALEHESYEMGEARFRKMFERQLKAGEVADNAAAKPLITTLLPKMIARINDWFEEVKAKRGK<br>RPTAFQFLQEIKPEAVAYITIKTTLACLTSADNTTVQAVASAIGRAIEDEARFGRIRDLEAKHFKKNVEEQLNKRVGHVY<br>KKAFMQVVEADMLSKGLLGGEAWSSWHKEDSIHVGVRCIEMLIESTGMVSLHRQNAGVVGQDSETIELAPEYAEAIATRA<br>GALAGISPMFQPCVVPPKPWTGITGGGYWANGRRPLALVRTHSKKALMRYEDVYMPEVYKAINIAQNTAWKINKKVLAVA<br>NVITKWKHCPVEDIPAIEREELPMKPEDIDMNPEALTAWKRAAAAVYRKDKARKSRRISLEFMLEQANKFANHKAIWFPY<br>NMDWRGRVYAVSMFNPQGNDMTKGLLTLAKGKPIGKEGYYWLKIHGANCAGVDKVPFPERIKFIEENHENIMACAKSPLE<br>NTWWAEQDSPFCFLAFCFEYAGVQHHGLSYNCSLPLAFDGSCSGIQHFSAMLRDEVGGRAVNLLPSETVQDIYGIVAKKV<br>NEILQADAINGTDNEVVTVTDENTGEISEKVKLGTKALAGQWLAYGVTRSVTKRSVMTLAYGSKEFGFRQQVLEDTIQPA<br>IDSGKGLMFTQPNQAAGYMAKLIWESVSVTVVAAVEAMNWLKSAAKLLAAEVKDKKTGEILRKRCAVHWVTPDGFPVWQE<br>YKKPIQTRLNLMFLGQFRLQPTINTNKDSEIDAHKQESGIAPNFVHSQDGSHLRKTVVWAHEKYGIESFALIHDSFGTIP<br>ADAANLFKAVRETMVDTYESCDVLADFYDQFADQLHESQLDKMPALPAKGNLNLRDILESDFAFA |
| pmCDA1-T7RNAP | MKRTADGSEFESPKKKRKVSTDAEYVRIHEKLDIYTFKKQFFNNKKSVSHRCYVLFELKRRGERRACFWGYAVNKPQSGT<br>ERGIHAEIFSIRKVEEYLRDNPGQFTINWYSSWSPCADCAEKILEWYNQELRGNGHTLKIWACKLYYEKNARNQIGLWNL<br>RDNGVGLNVMVSEHYQCCRKIFIQSSHNQLNENRWLEKTLKRAEKRRSELSIMIQVKILHTTKSPAVSGGSSGGSSGGGS<br>ETPGTSESATPESGGSDYKDDDDKGSLIKGMNTINIAKNEFLEPPKKKRKVEFSDIELAAIPFNTLADHYGERLAREQLA<br>LEHESYEMGEARFRKMFERQLKAGEVADNAAAKPLITTLLPKMIARINDWFEEVKAKRGKRPTAFQFLQEIKPEAVAYIT<br>IKTTLACLTSADNTTVQAVASAIGRAIEDEARFGRIRDLEAKHFKKNVEEQLNKRVGHVYKKAFMQVVEADMLSKGLLGG<br>EAWSSWHKEDSIHVGVRCIEMLIESTGMVSLHRQNAGVVGQDSETIELAPEYAEAIATRAGALAGISPMFQPCVVPPKPW<br>TGITGGGYWANGRRPLALVRTHSKKALMRYEDVYMPEVYKAINIAQNTAWKINKKVLAVANVITKWKHCPVEDIPAIERE<br>ELPMKPEDIDMNPEALTAWKRAAAAVYRKDKARKSRRISLEFMLEQANKFANHKAIWFPYNMDWRGRVYAVSMFNPQGND<br>MTKGLLTLAKGKPIGKEGYYWLKIHGANCAGVDKVPFPERIKFIEENHENIMACAKSPLENTWWAEQDSPFCFLAFCFEY<br>AGVQHHGLSYNCSLPLAFDGSCSGIQHFSAMLRDEVGGRAVNLLPSETVQDIYGIVAKKVNEILQADAINGTDNEVVTVT<br>DENTGEISEKVKLGTKALAGQWLAYGVTRSVTKRSVMTLAYGSKEFGFRQQVLEDTIQPAIDSGKGLMFTQPNQAAGYMA<br>KLIWESVSVTVVAAVEAMNWLKSAAKLLAAEVKDKKTGEILRKRCAVHWVTPDGFPVWQEYKKPIQTRLNLMFLGQFRLQ<br>PTINTNKDSEIDAHKQESGIAPNFVHSQDGSHLRKTVVWAHEKYGIESFALIHDSFGTIPADAANLFKAVRETMVDTYES<br>CDVLADFYDQFADQLHESQLDKMPALPAKGNLNLRDILESDFAFA |
| evoAPO1-evoTadA-T7RNAP (YeastIT) | MKRTADGSEFESPKKKRKVSSKTGPVAVDPTLRRRIEPHEFEVFFDPRELRKETCLLYEINWGGRHSIWRHTSQNTNKHV<br>EVNFIEKFTTERYFCPNTRCSITWFLSWSPCGECSRAITEFLSRYPNVTLFIYIARLYHLANPRNRQGLRDLISSGVTIQ<br>IMTEQESGYCWHNFVNYSPSNESHWPRYPHLWVRLYVLELYCIILGLPPCLNILRRKQSQLTSFTIALQSCHYQRLPPHI<br>LWATGLKSGGGSETPGTSESATPESGGSSEVEFSHEYWMRHALTLAKRARDEREVPVGAVLVLNNRVIGEGWNRAIGLHD<br>PTAHAEIMALRQGGLVMQNYRLIDATLYVTFEPCVMCAGAMIHSRIGRVVFGVRNSKRGAAGSLMNVLNYPGMNHRVEIT<br>EGILADECAALLCDFYRMPRQVFNAQKKAQSSINSGGGSETPGTSESATPESGGSIKGMNTINIAKNEFLEPPKKKRKVE<br>FSDIELAAIPFNTLADHYGERLAREQLALEHESYEMGEARFRKMFERQLKAGEVADNAAAKPLITTLLPKMIARINDWFE<br>EVKAKRGKRPTAFQFLQEIKPEAVAYITIKTTLACLTSADNTTVQAVASAIGRAIEDEARFGRIRDLEAKHFKKNVEEQL<br>NKRVGHVYKKAFMQVVEADMLSKGLLGGEAWSSWHKEDSIHVGVRCIEMLIESTGMVSLHRQNAGVVGQDSETIELAPEY<br>AEAIATRAGALAGISPMFQPCVVPPKPWTGITGGGYWANGRRPLALVRTHSKKALMRYEDVYMPEVYKAINIAQNTAWKI<br>NKKVLAVANVITKWKHCPVEDIPAIEREELPMKPEDIDMNPEALTAWKRAAAAVYRKDKARKSRRISLEFMLEQANKFAN<br>HKAIWFPYNMDWRGRVYAVSMFNPQGNDMTKGLLTLAKGKPIGKEGYYWLKIHGANCAGVDKVPFPERIKFIEENHENIM<br>ACAKSPLENTWWAEQDSPFCFLAFCFEYAGVQHHGLSYNCSLPLAFDGSCSGIQHFSAMLRDEVGGRAVNLLPSETVQDI<br>YGIVAKKVNEILQADAINGTDNEVVTVTDENTGEISEKVKLGTKALAGQWLAYGVTRSVTKRSVMTLAYGSKEFGFRQQV<br>LEDTIQPAIDSGKGLMFTQPNQAAGYMAKLIWESVSVTVVAAVEAMNWLKSAAKLLAAEVKDKKTGEILRKRCAVHWVTP<br>DGFPVWQEYKKPIQTRLNLMFLGQFRLQPTINTNKDSEIDAHKQESGIAPNFVHSQDGSHLRKTVVWAHEKYGIESFALI<br>HDSFGTIPADAANLFKAVRETMVDTYESCDVLADFYDQFADQLHESQLDKMPALPAKGNLNLRDILESDFAFA |

Domains are coloured by identity: cytidine deaminase (light green), adenosine deaminase (dark green), linker sequence (blue), SV40 nuclear localization signal (pink), FLAG tag (orange), T7 RNA polymerase (yellow, introduced point mutations are underlined), and P2A peptide (light blue).

### Supplementary Table 4. Mutator protein sequences used in this study (continued)

| | |
|---|---|
| pmCDA1-evoTadA-T7RNAP | MKRTADGSEFESPKKKRKVSTDAEYVRIHEKLDIYTFKKQFFNNKKSVSHRCYVLFELKRRGERRACFWGYAVNKPQSGT<br>ERGIHAEIFSIRKVEEYLRDNPGQFTINWYSSWSPCADCAEKILEWYNQELRGNGHTLKIWACKLYYEKNARNQIGLWNL<br>RDNGVGLNVMVSEHYQCCRKIFIQSSHNQLNENRWLEKTLKRAEKRRSELSIMIQVKILHTTKSPAVSGGGSETPGTSES<br>ATPESGGSSEVEFSHEYWMRHALTLAKRARDEREVPVGAVLVLNNRVIGEGWNRAIGLHDPTAHAEIMALRQGGLVMQNY<br>RLIDATLYVTFEPCVMCAGAMIHSRIGRVVFGVRNSKRGAAGSLMNVLNYPGMNHRVEITEGILADECAALLCDFYRMPR<br>QVFNAQKKAQSSINSGGGSETPGTSESATPESGGSIKGMNTINIAKNEFLEPPKKKRKVEFSDIELAAIPFNTLADHYGE<br>RLAREQLALEHESYEMGEARFRKMFERQLKAGEVADNAAAKPLITTLLPKMIARINDWFEEVKAKRGKRPTAFQFLQEIK<br>PEAVAYITIKTTLACLTSADNTTVQAVASAIGRAIEDEARFGRIRDLEAKHFKKNVEEQLNKRVGHVYKKAFMQVVEADM<br>LSKGLLGGEAWSSWHKEDSIHVGVRCIEMLIESTGMVSLHRQNAGVVGQDSETIELAPEYAEAIATRAGALAGISPMFQP<br>CVVPPKPWTGITGGGYWANGRRPLALVRTHSKKALMRYEDVYMPEVYKAINIAQNTAWKINKKVLAVANVITKWKHCPVE<br>DIPAIEREELPMKPEDIDMNPEALTAWKRAAAAVYRKDKARKSRRISLEFMLEQANKFANHKAIWFPYNMDWRGRVYAVS<br>MFNPQGNDMTKGLLTLAKGKPIGKEGYYWLKIHGANCAGVDKVPFPERIKFIEENHENIMACAKSPLENTWWAEQDSPFC<br>FLAFCFEYAGVQHHGLSYNCSLPLAFDGSCSGIQHFSAMLRDEVGGRAVNLLPSETVQDIYGIVAKKVNEILQADAINGT<br>DNEVVTVTDENTGEISEKVKLGTKALAGQWLAYGVTRSVTKRSVMTLAYGSKEFGFRQQVLEDTIQPAIDSGKGLMFTQP<br>NQAAGYMAKLIWESVSVTVVAAVEAMNWLKSAAKLLAAEVKDKKTGEILRKRCAVHWVTPDGFPVWQEYKKPIQTRLNLM<br>FLGQFRLQPTINTNKDSEIDAHKQESGIAPNFVHSQDGSHLRKTVVWAHEKYGIESFALIHDSFGTIPADAANLFKAVRE<br>TMVDTYESCDVLADFYDQFADQLHESQLDKMPALPAKGNLNLRDILESDFAFA |
| pmCDA1-T7RNAP* | MKRTADGSEFESPKKKRKVSTDAEYVRIHEKLDIYTFKKQFFNNKKSVSHRCYVLFELKRRGERRACFWGYAVNKPQSGT<br>ERGIHAEIFSIRKVEEYLRDNPGQFTINWYSSWSPCADCAEKILEWYNQELRGNGHTLKIWACKLYYEKNARNQIGLWNL<br>RDNGVGLNVMVSEHYQCCRKIFIQSSHNQLNENRWLEKTLKRAEKRRSELSIMIQVKILHTTKSPAVSGGSSGGSSGGGS<br>ETPGTSESATPESGGSDYKDDDDKGSLIKGMNTINIAKNEFLEPPKKKRKVEFSDIELAAIPFNTLADHYGERLAREQLA<br>LEHESYEMGEARFRKMFERQLKAGEVADNAAAKPLITTLLPKMIARINDWFEEVKAKRGKRPTAFQFLQEIKPEAVAYIT<br>IKTTLACLTSADNTTVQAVASAIGRAIEDEARFGRIRDLEAKHFKKNVEEQLNKRVGHVYKKAFMQVVEADMLSKGLLGG<br>EAWSSWHKEDSIHVGVRCIEMLIESTGMVSLHRQNAGVVGQDSETIELAPEYAEAIATRAGALAGISPMFQPCVVPPKPW<br>TGITGGGYWANGRRPLALVRTHSKKALMRYEDVYMPEVYKAINIAQNTAWKINKKVLAVANVITKWKHCPVEDIPAIERE<br>ELPMKPEDIDMNPEALTAWKRAAAAVYRKDKARKSRRISLEFMLEQANKFANHKAIWFPYNMDWRGRVYAVSMFNPQGND<br>MTKGLLTLAKGKPIGKEGYYWLKIHGANCAGVDKVPFPERIKFIEENHENIMACAKSPLENTWWAEQDSPFCFLAFCFEY<br>AGVQHHGLSYNCSLPLAFDGSCSGIQHFSAMLRDEVGGRAVNLLPSETVQDIYGIVAKKVNEILQADAINGTDNEVVTVT<br>DENTGEISEKVKLGTKALAGQWLAYGVTRSVTKRSVMTLAYGSKEF**A**FRQQVLEDTIQPAIDSGKGLMFTQPNQAAGYMA<br>KLIWESVSVTVVAAVEAMNWLKSAAKLLAAEVKDKKTGEILRKRCAVHWVTPDGFPVWQEYKKPI**R**TRLNLMFLGQFRLQ<br>PTINTNKDSEIDAHKQESGIAPNFVHSQDGSHLRKTVVWAHEKYGIESFALIHDSFGTIPADAANLFKAVRETMVDTYES<br>CDVLADFYDQFADQLHESQLDKMPALPAKGNLNLRDILESDFAFA |
| pmCDA1-29aa-T7RNAP | MKRTADGSEFESPKKKRKVSTDAEYVRIHEKLDIYTFKKQFFNNKKSVSHRCYVLFELKRRGERRACFWGYAVNKPQSGT<br>ERGIHAEIFSIRKVEEYLRDNPGQFTINWYSSWSPCADCAEKILEWYNQELRGNGHTLKIWACKLYYEKNARNQIGLWNL<br>RDNGVGLNVMVSEHYQCCRKIFIQSSHNQLNENRWLEKTLKRAEKRRSELSIMIQVKILHTTKSPAVGGSETPGTSESAT<br>PESGGSDYKDDDDKGSLIKGMNTINIAKNEFLEPPKKKRKVEFSDIELAAIPFNTLADHYGERLAREQLALEHESYEMGE<br>ARFRKMFERQLKAGEVADNAAAKPLITTLLPKMIARINDWFEEVKAKRGKRPTAFQFLQEIKPEAVAYITIKTTLACLTS<br>ADNTTVQAVASAIGRAIEDEARFGRIRDLEAKHFKKNVEEQLNKRVGHVYKKAFMQVVEADMLSKGLLGGEAWSSWHKED<br>SIHVGVRCIEMLIESTGMVSLHRQNAGVVGQDSETIELAPEYAEAIATRAGALAGISPMFQPCVVPPKPWTGITGGGYWA<br>NGRRPLALVRTHSKKALMRYEDVYMPEVYKAINIAQNTAWKINKKVLAVANVITKWKHCPVEDIPAIEREELPMKPEDID<br>MNPEALTAWKRAAAAVYRKDKARKSRRISLEFMLEQANKFANHKAIWFPYNMDWRGRVYAVSMFNPQGNDMTKGLLTLAK<br>GKPIGKEGYYWLKIHGANCAGVDKVPFPERIKFIEENHENIMACAKSPLENTWWAEQDSPFCFLAFCFEYAGVQHHGLSY<br>NCSLPLAFDGSCSGIQHFSAMLRDEVGGRAVNLLPSETVQDIYGIVAKKVNEILQADAINGTDNEVVTVTDENTGEISEK<br>VKLGTKALAGQWLAYGVTRSVTKRSVMTLAYGSKEFGFRQQVLEDTIQPAIDSGKGLMFTQPNQAAGYMAKLIWESVSVT<br>VVAAVEAMNWLKSAAKLLAAEVKDKKTGEILRKRCAVHWVTPDGFPVWQEYKKPIQTRLNLMFLGQFRLQPTINTNKDSE<br>IDAHKQESGIAPNFVHSQDGSHLRKTVVWAHEKYGIESFALIHDSFGTIPADAANLFKAVRETMVDTYESCDVLADFYDQ<br>FADQLHESQLDKMPALPAKGNLNLRDILESDFAFA |
| pmCDA1-51aa-T7RNAP | MKRTADGSEFESPKKKRKVSTDAEYVRIHEKLDIYTFKKQFFNNKKSVSHRCYVLFELKRRGERRACFWGYAVNKPQSGT<br>ERGIHAEIFSIRKVEEYLRDNPGQFTINWYSSWSPCADCAEKILEWYNQELRGNGHTLKIWACKLYYEKNARNQIGLWNL<br>RDNGVGLNVMVSEHYQCCRKIFIQSSHNQLNENRWLEKTLKRAEKRRSELSIMIQVKILHTTKSPAVSGGSSGGSSGGGS<br>ETPGTSESATPESGGSDYKDDDDKGSLISGGSSGGIKGMNTINIAKNEFLEPPKKKRKVEFSDIELAAIPFNTLADHYGE<br>RLAREQLALEHESYEMGEARFRKMFERQLKAGEVADNAAAKPLITTLLPKMIARINDWFEEVKAKRGKRPTAFQFLQEIK<br>PEAVAYITIKTTLACLTSADNTTVQAVASAIGRAIEDEARFGRIRDLEAKHFKKNVEEQLNKRVGHVYKKAFMQVVEADM<br>LSKGLLGGEAWSSWHKEDSIHVGVRCIEMLIESTGMVSLHRQNAGVVGQDSETIELAPEYAEAIATRAGALAGISPMFQP<br>CVVPPKPWTGITGGGYWANGRRPLALVRTHSKKALMRYEDVYMPEVYKAINIAQNTAWKINKKVLAVANVITKWKHCPVE<br>DIPAIEREELPMKPEDIDMNPEALTAWKRAAAAVYRKDKARKSRRISLEFMLEQANKFANHKAIWFPYNMDWRGRVYAVS<br>MFNPQGNDMTKGLLTLAKGKPIGKEGYYWLKIHGANCAGVDKVPFPERIKFIEENHENIMACAKSPLENTWWAEQDSPFC<br>FLAFCFEYAGVQHHGLSYNCSLPLAFDGSCSGIQHFSAMLRDEVGGRAVNLLPSETVQDIYGIVAKKVNEILQADAINGT<br>DNEVVTVTDENTGEISEKVKLGTKALAGQWLAYGVTRSVTKRSVMTLAYGSKEFGFRQQVLEDTIQPAIDSGKGLMFTQP<br>NQAAGYMAKLIWESVSVTVVAAVEAMNWLKSAAKLLAAEVKDKKTGEILRKRCAVHWVTPDGFPVWQEYKKPIQTRLNLM |

| | |
|---|---|
| | FLGQFRLQPTINTNKDSEIDAHKQESGIAPNFVHSQDGSHLRKTVVWAHEKYGIESFALIHDSFGTIPADAANLFKAVRE<br>TMVDTYESCDVLADFYDQFADQLHESQLDKMPALPAKGNLNLRDILESDFAFA |

**Supplementary Table 4. Mutator protein sequences used in this study (continued)**

| | |
|---|---|
| pmCDA1-P2A-RNAP | MKRTADGSEFESPKKKRKVSTDAEYVRIHEKLDIYTFKKQFFNNKKSVSHRCYVLFELKRRGERRACFWGYAVNKPQSGT<br>ERGIHAEIFSIRKVEEYLRDNPGQFTINWYSSWSPCADCAEKILEWYNQELRGNGHTLKIWACKLYYEKNARNQIGLWNL<br>RDNGVGLNVMVSEHYQCCRKIFIQSSHNQLNENRWLEKTLKRAEKRRSELSIMIQVKILHTTKSPAVSGGSSGGSSGGGS<br>ETPGTSESATPESGGSDYKDDDDKGSLIGSGATNFSLLKQAGDVEENPGPGSIKGMNTINIAKNEFLEPPKKKRKVEFSD<br>IELAAIPFNTLADHYGERLAREQLALEHESYEMGEARFRKMFERQLKAGEVADNAAAKPLITTLLPKMIARINDWFEEVK<br>AKRGKRPTAFQFLQEIKPEAVAYITIKTTLACLTSADNTTVQAVASAIGRAIEDEARFGRIRDLEAKHFKKNVEEQLNKR<br>VGHVYKKAFMQVVEADMLSKGLLGGEAWSSWHKEDSIHVGVRCIEMLIESTGMVSLHRQNAGVVGQDSETIELAPEYAEA<br>IATRAGALAGISPMFQPCVVPPKPWTGITGGGYWANGRRPLALVRTHSKKALMRYEDVYMPEVYKAINIAQNTAWKINKK<br>VLAVANVITKWKHCPVEDIPAIEREELPMKPEDIDMNPEALTAWKRAAAAVYRKDKARKSRRISLEFMLEQANKFANHKA<br>IWFPYNMDWRGRVYAVSMFNPQGNDMTKGLLTLAKGKPIGKEGYYWLKIHGANCAGVDKVPFPERIKFIEENHENIMACA<br>KSPLENTWWAEQDSPFCFLAFCFEYAGVQHHGLSYNCSLPLAFDGSCSGIQHFSAMLRDEVGGRAVNLLPSETVQDIYGI<br>VAKKVNEILQADAINGTDNEVVTVTDENTGEISEKVKLGTKALAGQWLAYGVTRSVTKRSVMTLAYGSKEFGFRQQVLED<br>TIQPAIDSGKGLMFTQPNQAAGYMAKLIWESVSVTVVAAVEAMNWLKSAAKLLAAEVKDKKTGEILRKRCAVHWVTPDGF<br>PVWQEYKKPIQTRLNLMFLGQFRLQPTINTNKDSEIDAHKQESGIAPNFVHSQDGSHLRKTVVWAHEKYGIESFALIHDS<br>FGTIPADAANLFKAVRETMVDTYESCDVLADFYDQFADQLHESQLDKMPALPAKGNLNLRDILESDFAFA |

**Supplementary Table 5. Mutagenesis target sequences used in this study**

| Name | Sequence[a] |
|---|---|
| she1 | MNDKLQEEHNEKDTTSQINGFTPPHMSIDFHSNNNSNIIETIGVSKRLGNSVLSELDSRASSKFEFLKDQSEQQYNGDKN<br>NEPKSGSYNINEFFQAKHDSQFGQMESLDTHYTLLHTPKRKSQHAIPQDRSDSMKRSRPSRSIPYTTPVVNDITRRIRRL<br>KLRNSLVNGNDIVARARSMQANSNINSIKNTPLSKPKPFMHKPNFLMPTTNSLNKINSAHRNTSSSSTASSIPRSKVHRS<br>ISIRDLHAKTKPVERTPVAQGTNSQLKNSVSVFDRLYKQTTFSRSTSMNNLSSGTSAKSKEHTNVKTRLVKSKTSGSLSS<br>NLKQSTATGTKSDRPIWR |
| Aga2-3G146 | MQLLRCFSIFSVIASVLAQELTTICEQIPSPTLESTPYSLSTTTILANGKAMQGVFEYYKSVTFVSNCGSHPSTTSKGSP<br>INTQYVFKDNSSTIEGRYPYDVPDYALQASGGGGSGGGGSGGGGSDLGKKLLEAARAGQDDEVRILMANGADVNAIDQFG<br>DTPLHLAASFGHLEIVEVLLKYGVDVNARDVAGLTPLHLAAKWGHFEIVEVLLKHGADVNASDVWGWTPLHLAAAMGHLE<br>IVEVLLKYGADVNAQDKFGKTAFDISINNGNEDLAEILQKLNGSEGKLISEEDL |

[a] Domains are coloured by identity: aga2 signal peptide (light blue), aga2 (yellow), linker sequence (blue), HA tag (orange), 3G146 (pink), c-myc tag (green)

**Supplementary Table 6. Protein sequences of *E.* coli-expressed DARPin variants**

| Name | Sequence[a] |
|---|---|
| 3G146 | MYPYDVPDYALQASGGGGSGGGGSGGGGSDLGKKLLEAARAGQDDEVRILMANGADVNAIDQFGDTPLHLAASFGHLEIV<br>EVLLKYGVDVNARDVAGLTPLHLAAKWGHFEIVEVLLKHGADVNASDVWGWTPLHLAAAMGHLEIVEVLLKYGADVNAQD<br>KFGKTAFDISINNGNEDLAEILQKLNMGLGSHHHHHH |
| 2-G4 | MYPYDVPDYALQASGGGGSGGGGSGGGGSDLGKKLLEAARAGQDDEVRILMAN**S**ADVNAIDQFGDTPLHLAASFGHLEIV<br>EVLLKYGVDVNARDVAGLTPLHLAAKWGHFEIVEVLLKHGADVNASDVWGWTPLHLAAAMGHLEIVEVLLKYGADVNAQD<br>KFGKTAFDISINNGNEDLAEILQKLNMGLGSHHHHHH |
| 2-H3 | MYPYDVPDYALQASGGGGSGGGGSGGGGSDLGKKLLEAARAGQDDEVRILMANGADVNAIDQFGDTPLHLAASFGHLEIV<br>EVLLKYGVDVNARDVAGLTPLHLAAKWGHFEIVEVLLKHGADVNASDVWGWTPLHLAAAM**S**HLEIVEVLLKYGADVNAQD<br>KFGKTAFDISINNGNEDLAEILQKLNMGLGSHHHHHH |

[a] Domains are coloured by identity: linker sequence (blue), HA tag (orange), 3G146 (pink), $His_6$-tag (green). Mutations in discovered variants are indicated in bold.

### Supplementary Table 7. Mutational spectra of random mutagenesis methods

| Method | Mutation spectrum (%) | | | | | | Reference |
|---|---|---|---|---|---|---|---|
| | AT>GC | GC>AT | AT>CG | AT>TA | GC>TA | GC>CG | |
| YeastIT1 | 18.32 | 81.12 | 0.08 | 0 | 0.48 | 0 | This work |
| YeastIT2 | 77.11 | 5.97 | 0.50 | 1 | 4.98 | 10.45 | This work |
| MutaT7 | 0 | 100 | 0 | 0 | 0 | 0 | Moore et al.[2], Figure S10b |
| eMutaT7 | 0 | 100 | 0 | 0 | 0 | 0 | Park et al.[3], Figure 3a |
| EvolVR | 25.32 | 29.54 | 25.32 | 14.77 | 3.38 | 1.69 | Halperin et al.[4], Figure 3c |
| PACE | 12.63 | 45.26 | 8.42 | 10.53 | 15.79 | 7.37 | Esvelt et al.[5], Figure S5 |
| OrthoRep | 73.13 | 21.33 | 1.1 | 1.1 | 1.1 | 2.23 | Ravikumar et al.[6], Table S8 |
| TRIDENT | 3 | 97 | 0 | 0 | 0 | 0 | Cravens et al., reanalysed[a] |
| epPCR (Taq, Mn2+) | 64.82 | 9.75 | 4.1 | 16.51 | 3.38 | 1.43 | Wong et al.[7], Table 1 |
| epPCR (Mutazyme II) | 18.54 | 27.01 | 4.98 | 30.19 | 14.93 | 4.34 | Wong et al.[7], Table 1 |

[a] Raw data were accessed at NCBI (accession number: PRJNA701053). Analysis was done using the scripts provided by the authors[8]. In order to allow for a more accurate comparison with other methods, the window of analysis was altered from 200 bp to a full-length gene. The analysis can be accessed at: https://github.com/fhlab/YeastIT/tree/main/TRIDENT-reanalysis

**Supplementary Table 8. Overview of point mutations revealed by Sanger sequencing of *she1* mutants generated by different mutator variants.** Mutations were observed in clones retrieved from experiments both with and without selection pressure. No insertions or deletions were present in the sequenced variants.

| Mutator variant | Selection pressure | Mutations in *she1* CDS | No. of mutations | She1 aa substitutions |
|---|---|---|---|---|
| Δung1 evoAPO1 | yes | 49C>T | 1 | Q17X |
| Δung1 evoAPO1 | yes | 49C>T | 1 | Q17X |
| Δung1 evoAPO1 | yes | 95C>T, 689C>T | 2 | S32F, S231F |
| Δung1 pmCDA1 | yes | 49C>T, 162C>T | 2 | Q17X |
| Δung1 pmCDA1 | yes | 630C>T, 793C>T | 2 | Q265X |
| Δung1 pmCDA1 | yes | 49C>T, 793C>T | 2 | Q17X, Q265X |
| Δung1 pmCDA1 | yes | 49C>T | 1 | Q17X |
| Δung1 pmCDA1 | no | 843C>T | 1 | - |
| Δung1 pmCDA1 | no | 674C>T | 1 | S225L |
| YeastIT1 | yes | 208C>T | 1 | Q70X |
| YeastIT1 | yes | 42C>T, 80C>T, 123C>T, 161C>T | 4 | S27F, S54F |
| YeastIT1 | yes | 108C>T, 208C>T, 307G>A, 366C>T, 419C>T, 430C>T, 505G>A, 548C>T | 8 | Q70X, G103S, S140F, P144S, G169S, S183F |
| YeastIT1 | yes | 42C>T, 45C>T, 49C>T, 73C>T, 80C>T, 84C>T, 162C>T, 201G>A, 366C>T, 367C>T, 381A>G, 431C>T, 675C>T, 697C>T | 14 | Q17X, H25Y, S27F, Q123X, P144L, S225L, P233S |
| YeastIT2 | no | 35A>G, 234T>C | 2 | K12R |
| YeastIT2 | no | 39T>C | 2 | K136E |
| YeastIT2 | no | 136A>G | 1 | K46E |

**Supplementary Table 9. Distribution of non-synonymous mutations in a random mutagenesis library of a 338 amino acid-long protein sequence.** Mutation frequencies ε were derived from in YeastIT1 and YeastIT2 UMIC-seq experiments. $k$ – number of substitutions at the amino acid level, $D_{max}$ – maximum theoretical degeneracy of a library, $P_k$ – probability of having $k$ amino acid substitutions, $T_k$ – number of library members containing $k$ mutations for sampled library size of $T1=10^6$ or $T2=10^7$, $Ni$ – average frequency of each library variants with $k$ substitutions, $Nt$ – total number of unique library variants with $k$ substitutions. Calculations assume equal probabilities of each DNA mutation and at each position in the sequence are assumed and no two mutations occurring in the same codon.

| | | YeastIT1 (ε=0.00414) | | | | YeastIT2 (ε=0.00027) | | | | | | |
|---|---|---|---|---|---|---|---|---|---|---|---|---|
| $k$ | $D_{max}$ | $P_k$ | $T1_k$ | $N_{i(T1)}$ | $N_{t(T1)}$ | $P_k$ | $T1_k$ | $N_{i(T1)}$ | $N_{t(T1)}$ | $T2_k$ | $N_{i(T2)}$ | $N_{t(T2)}$ |
| 0 | 1 | 0.05 | 52265 | $5.2\times10^{4}$ | 1 | 0.83 | 825554 | $8.3\times10^{5}$ | 1 | 8255542 | $8.3\times10^{6}$ | 1 |
| 1 | $2.1\times10^{3}$ | 0.15 | 154933 | 72.8 | 2129 | 0.16 | 158304 | $7.4\times10^{1}$ | 2129 | 1583039 | $7.4\times10^{2}$ | 2129 |
| 2 | $2.3\times10^{6}$ | 0.23 | 228957 | $1.0\times10^{-1}$ | 217744 | 0.02 | 15133 | $6.7\times10^{-3}$ | 15082 | 151329 | $6.7\times10^{-2}$ | 146374 |
| 3 | $1.6\times10^{9}$ | 0.22 | 224898 | $1.4\times10^{-4}$ | 224882 | 0.00 | 962 | $6.0\times10^{-7}$ | 962 | 9615 | $6.0\times10^{-6}$ | 9615 |
| 4 | $8.4\times10^{11}$ | 0.17 | 165189 | $2.0\times10^{-7}$ | 165189 | 0.00 | 46 | $5.4\times10^{-11}$ | 46 | 457 | $5.4\times10^{-10}$ | 457 |
| 5 | $3.5\times10^{14}$ | 0.10 | 96776 | $2.7\times10^{-10}$ | 96776 | 0.00 | 2 | $4.9\times10^{-15}$ | 2 | 17 | $4.9\times10^{-14}$ | 17 |
| 6 | $1.2\times10^{17}$ | 0.05 | 47106 | $3.8\times10^{-13}$ | 47106 | 0.00 | 0 | $4.4\times10^{-19}$ | 0 | 1 | $4.4\times10^{-18}$ | 1 |
| 7 | $3.7\times10^{19}$ | 0.02 | 19594 | $5.3\times10^{-16}$ | 19594 | 0.00 | 0 | $4.0\times10^{-23}$ | 0 | 0 | $4.0\times10^{-22}$ | 0 |
| 8 | $9.6\times10^{21}$ | 0.01 | 7110 | $7.4\times10^{-19}$ | 7110 | 0.00 | 0 | $3.6\times10^{-27}$ | 0 | 0 | $3.6\times10^{-26}$ | 0 |
| 9 | $2.2\times10^{23}$ | 0.00 | 2286 | $1.0\times10^{-21}$ | 2286 | 0.00 | 0 | $3.2\times10^{-31}$ | 0 | 0 | $3.2\times10^{-30}$ | 0 |
| 10 | $4.6\times10^{26}$ | 0.00 | 660 | $1.4\times10^{-24}$ | 660 | 0.00 | 0 | $2.9\times10^{-35}$ | 0 | 0 | $2.9\times10^{-34}$ | 0 |

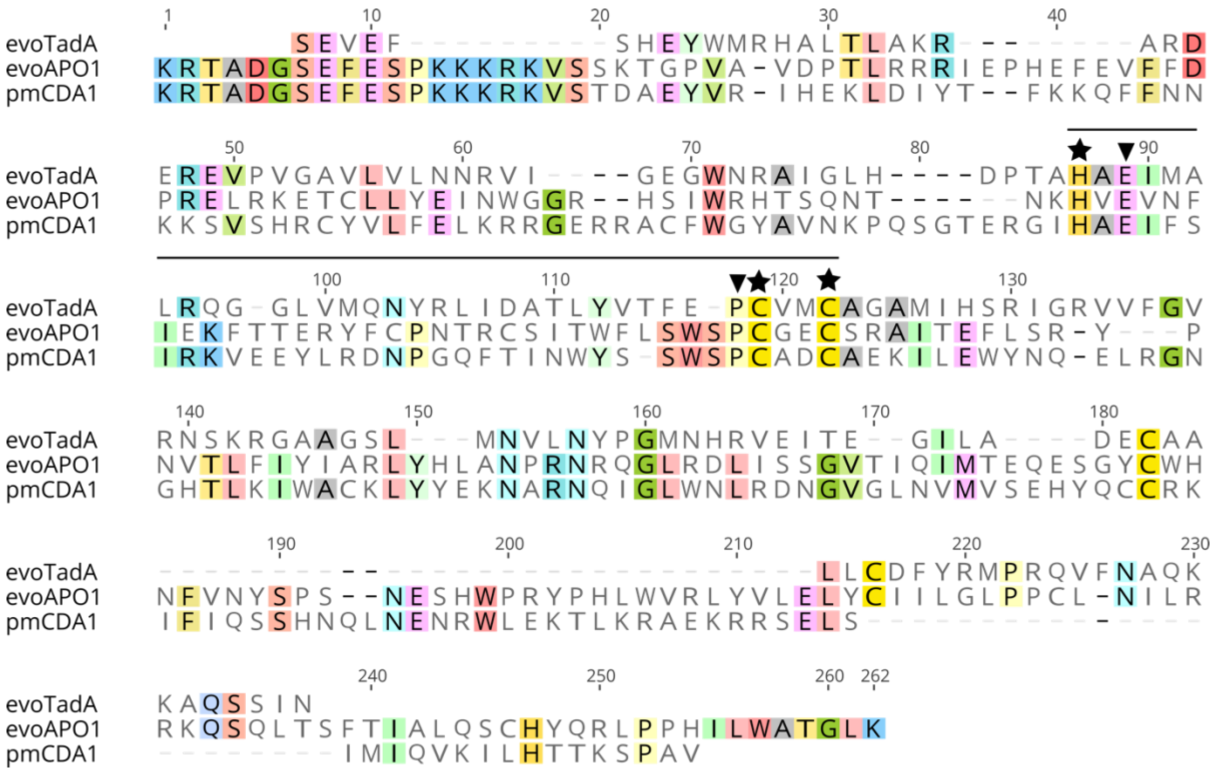

**Supplementary Figure 1. Multiple sequence alignment of three nucleoside deaminase modules used in this study: evoTadA, evoAPO1, and pmCDA1.** The conserved nucleoside deaminase domain H-x-E-x(24,28)-P-C-x(2,4)-C is necessary for catalytic activity[9]. Three $Zn^{2+}$-coordinating residues (★) and two catalytic residues (▼) are indicated. Sequence alignment was done with Geneious Prime using Clustal Omega algorithm, v. 1.2.0

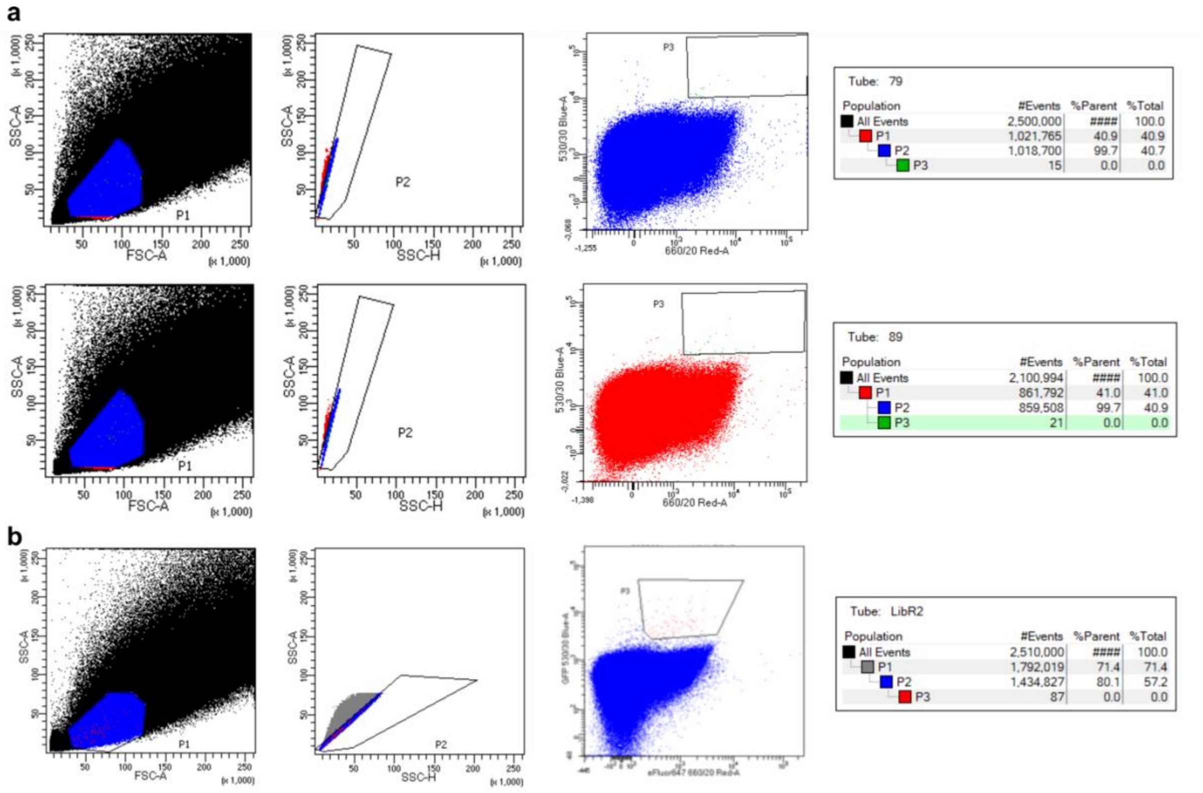

**Supplementary Figure 2. Gating strategy for sorting of YeastIT DARPin libraries.** P1 in the channels FSC area/SSC area and P2 in the channels SSC area/SSC height were used to exclude doublets and large cells. P3 in the channels EGFP (Ex 488 nm, Em 530/30 nm) area/iFluor 647 (Ex 633 nm, Em 660/20 nm) area was used to select cells with highest EGFP/iFluor 647 binding ratio. Less than 0.01% of cells were isolated in each round. (a) Gating in first round of directed evolution - YeastIT1 (upper diagrams) and YeastIT2 (lower diagrams) libraries were sorted separately. (b) Gating in second round of directed evolution (YeastIT1 and YeastIT2 libraries were combined prior to sorting).

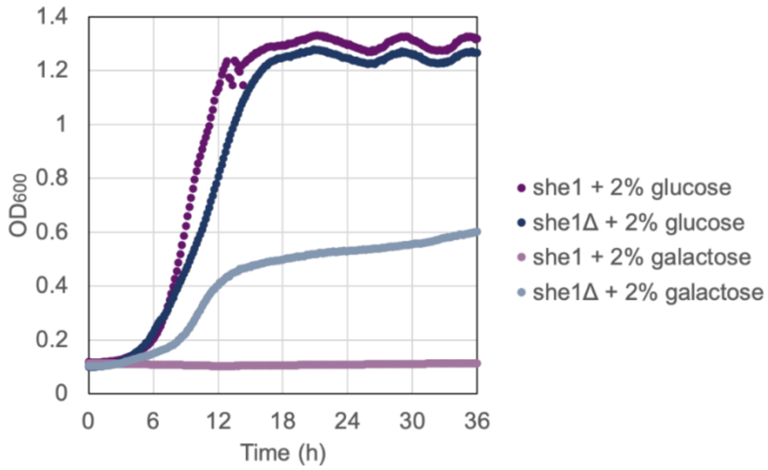

**Supplementary Figure 3. Growth curves of *S. cerevisiae* EBY100/pGoi-she1 and *S. cerevisiae* EBY100/pGoi-she1Δ in SDΔW and SGΔW.** *She1*, full-length gene sequence, *She1*Δ, inactive sequence that was C-terminally truncated by 92 amino acids. Strains were grown in SDΔW in deep-well plates until stationary phase, and 200 µl of fresh SDΔW or SGΔW was inoculated at starting OD of 0.1, transferred to 96-well U-bottom plates and sealed with transparent film. Cultures were grown for 36 hours at 30°C, shaking (4 mm amplitude) in Infinite 200 PRO microplate reader (Tecan) and absorbance (600 nm) was recorded every 10 min. Both strains exhibit equal growth rates in SDΔW. For EBY100/pGoi-she1Δ, lag-phase and reduced growth rate is observed when carbon source is changed to galactose, EBY100/pGoi-she1 does not grow on galactose-containing media.

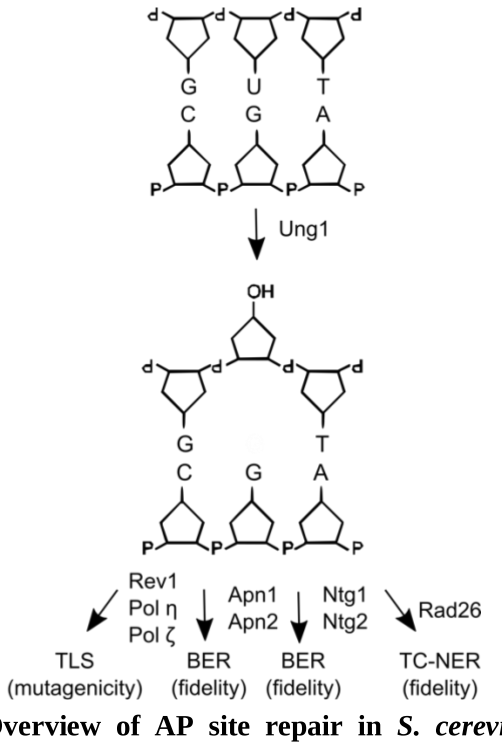

**Supplementary Figure 4. Overview of AP site repair in *S. cerevisiae*.** Excision of uracil by uracil-DNA glycosylase Ung1 generates an AP site. Error-free repair of generated AP sites is performed by base excision repair (BER) process initiated with the cleavage on the 5' side by Apn1 or Apn2, or with the cleavage on the 3' side by Ntg1 or Ntg2, or by transcription-coupled nucleotide excision repair (TC-NER), which can be initiated by Rad26[10,11]. Alternatively, AP site can be bypassed in the process of translesion synthesis (TLS), which can involve Rev1, Pol η and/or Pol ζ and is mostly mutagenic[11,12].

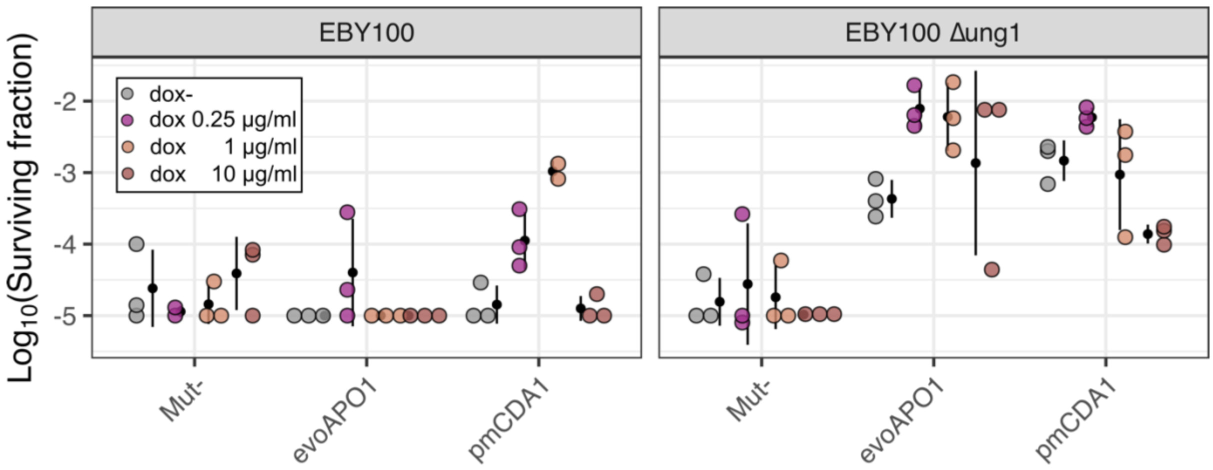

**Supplementary Figure 5. Mutation efficiency of mutator variants comprising of cytidine deaminase variants, evoAPO1 or pmCDA1, fused to T7RNAP at different concentrations of inducer (doxycycline).** Mutator performance was tested in different genetic backgrounds (EBY100 or EBY100 Δ*ung1*). Optimal efficiency is observed in *ung1*-deficient strain at 0.25 μg/ml of doxycycline with partial loss of mutator phenotype observed at higher doxycycline concentrations.

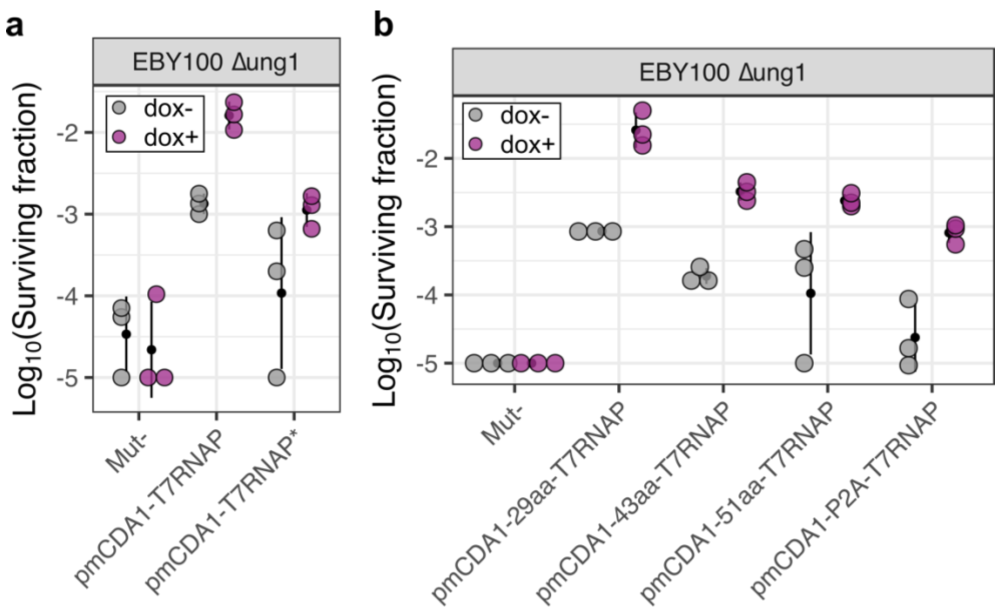

**Supplementary Figure 6. Mutation efficiency of pmCDA1-based mutators with different T7RNAP variants and varying length of linker sequences.** (**a**) Performance pmCDA1-T7RNAP and pmCDA1-T7RNAP*, a related variant containing 2 mutations (G645A, Q744R) in T7RNAP sequence. Introduction of mutations previously reported to be beneficial for mutagenic activity resulted in decreased mutation efficiency. (**b**) Performance of pmCDA1-T7RNAP variants with different linker lengths (sequences specified in **Supplementary Table 4**). Variant pmCDA1-43aa-T7RNAP is referred to as pmCDA1-T7RNAP in all other experiments. Increase in mutation efficiency with decreasing linker length is observed. P2A, 2A peptide that induces ribosomal skipping during translation.

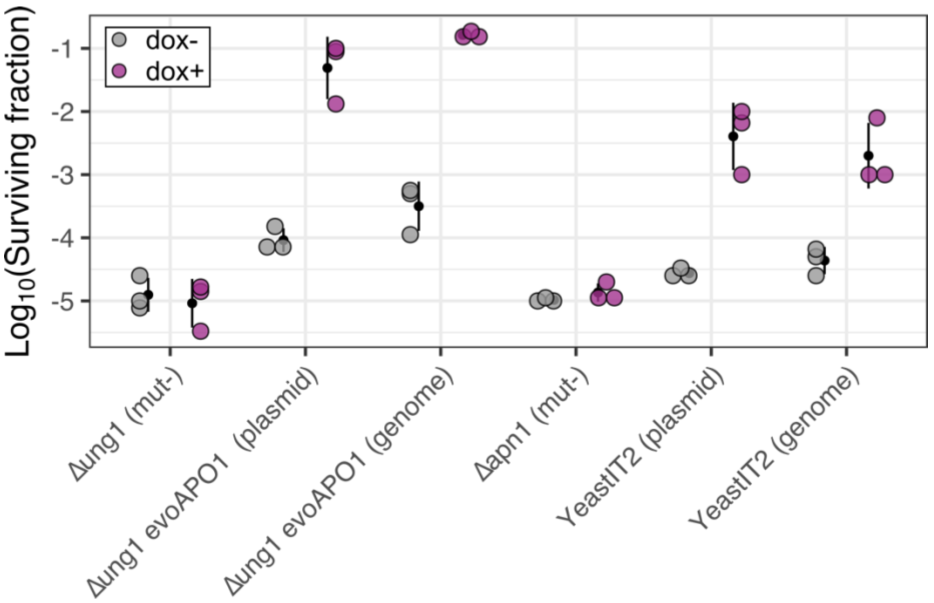

**Supplementary Figure 7. Mutation efficiency of evoAPO1-T7RNAP and YeastIT2 system in plasmid-based and genome-integrated versions.** Mutator phenotype was maintained when mutator proteins were integrated in the HO locus of yeast genome, but no improvement in mutation efficiency was observed.

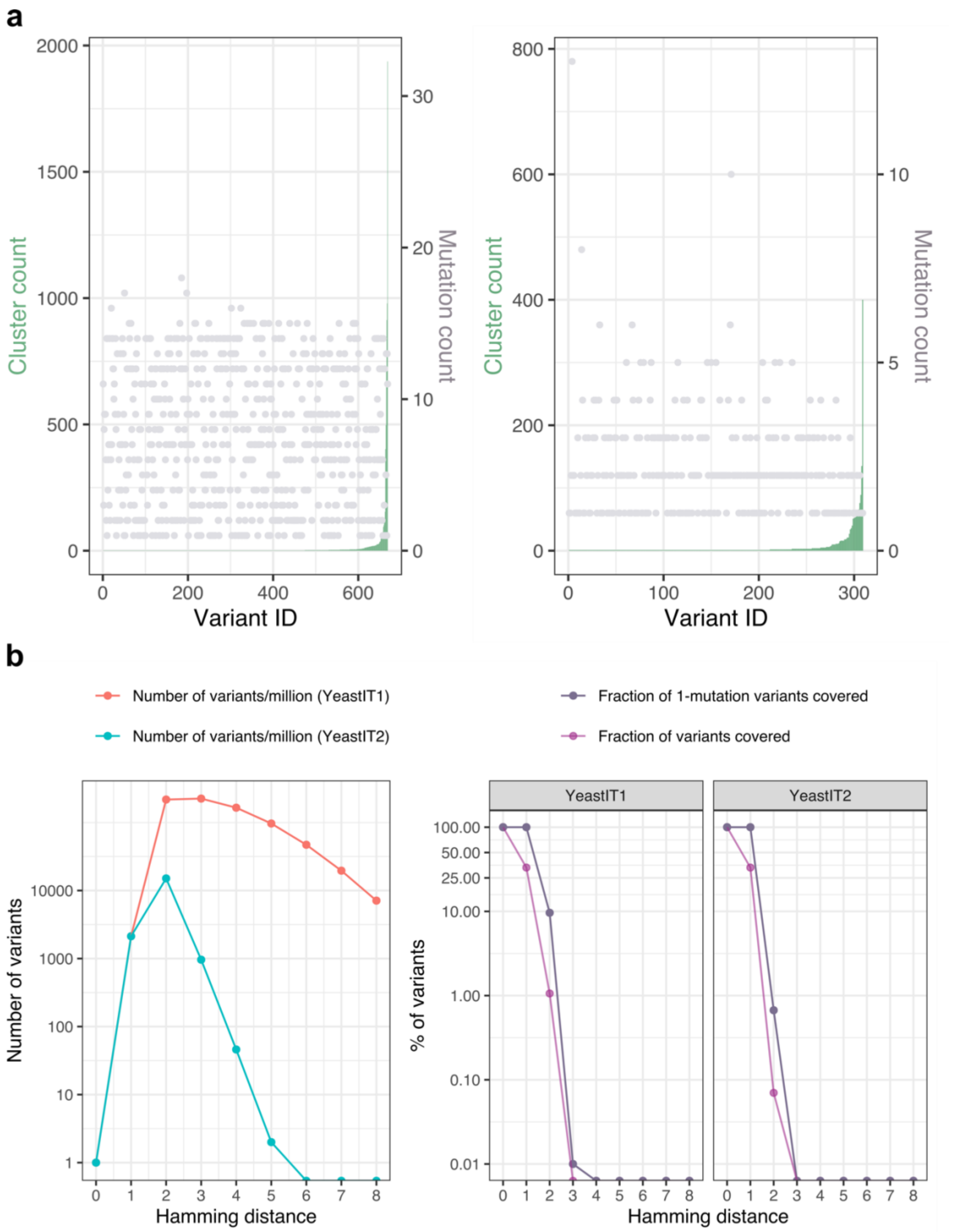

**Supplementary Figure 8. Estimation of total diversity in DNA libraries generated with YeastIT1 and YeastIT2. (a)** Distribution of UMI cluster counts and mutational counts in unique variants identified in UMIC-seq data. The mutational count shows uniform distribution across variants with different cluster abundance, indicating that the variants cannot be attributed to sequencing errors. 68% and 71% of variants were mapped to a single UMI cluster for YeastIT1 and YeastIT2, respectively, indicating that the true diversity of the generated DNA libraries is undersampled. **(b)** Estimation of library diversity per million cells achievable with YeastIT1 and YeastIT2 mutational rates as a function of Hamming distance to wild-type sequence displayed as the absolute number of unique variants (left) or as a percentage of the theoretical maximum diversity (right).

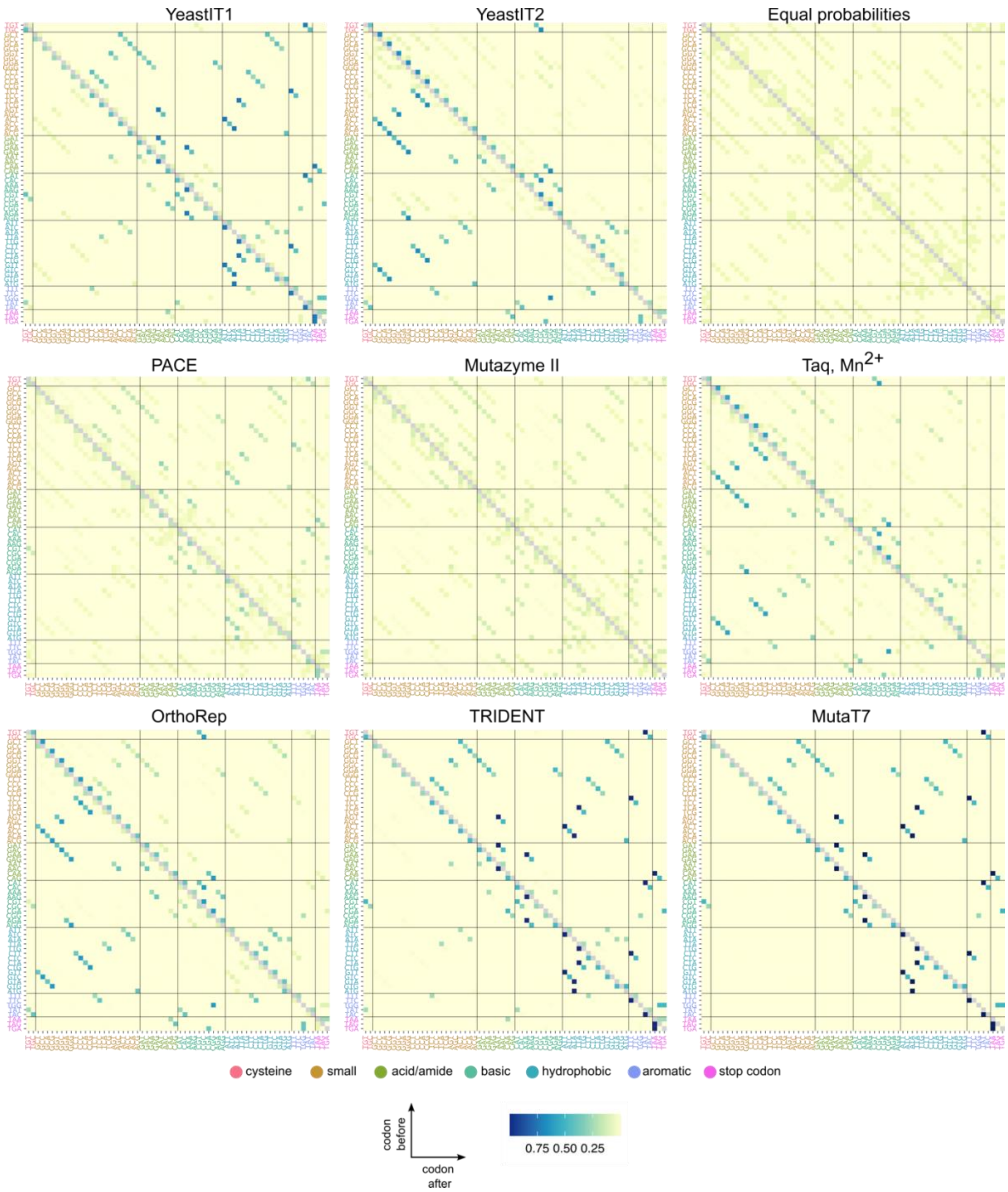

**Supplementary Figure 9. Codon substitution matrices of random mutagenesis methods and theoretical mutagenesis system performing every mutation with equal probability.** Matrices display the relative probability of each substitution for a given codon.

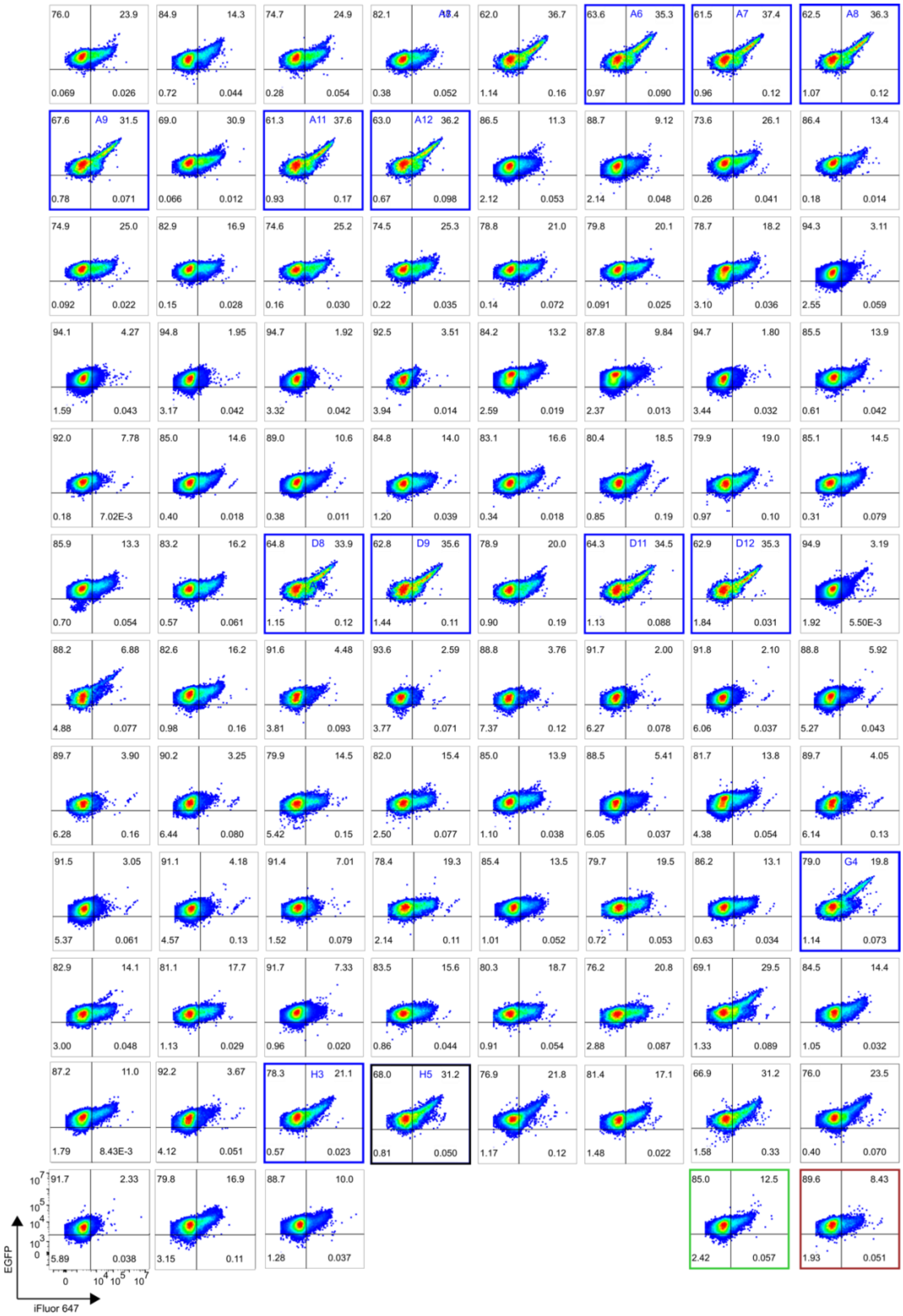

**Supplementary Figure 10. Flow cytometry data (N>10,000) of isogenic yeast cultures isolated after 2 rounds of directed evolution of 3G146-displaying *S. cerevisiae*.** Sorted cells were regrown on SDΔLW-agar plates, cultured in SGΔLW to display DARPin variants, stained with 1:1,000 iFluor 647 anti-HA antibody and 30 nM EGFP-$His_6$, and subjected to flow cytometry. Variants selected for Sanger sequencing are indicated in blue. Strain expressing wild-type 3G146 is indicated in green. 3G146-negative, HA tag-positive control is indicated in red. Percentage of cells in each quadrant (anti-HA+ EGFP+, anti-HA+, EGFP-, anti-HA- EGFP+, anti-HA- EGFP-) is indicated on the graphs.

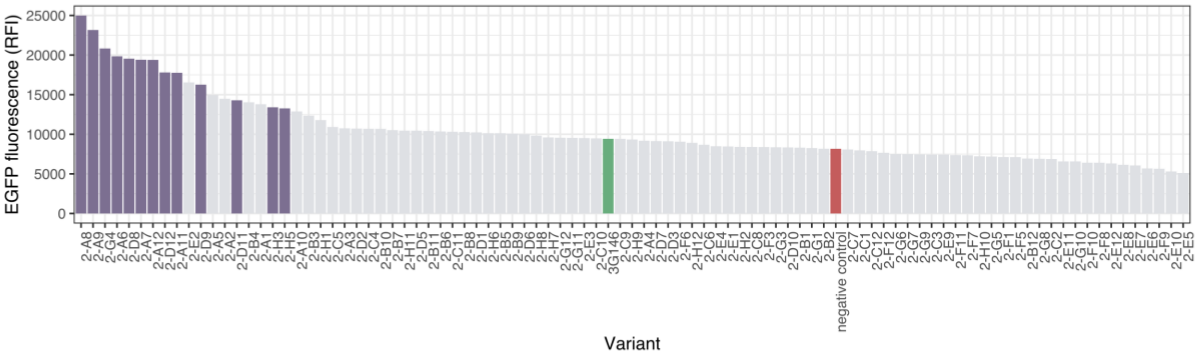

**Supplementary Figure 11. Summary of the median EGFP fluorescence values of display-positive fractions in isogenic yeast cultures isolated after 2 rounds of directed evolution of 3G146-displaying *S. cerevisiae***. Variants selected for Sanger sequencing are indicated in purple (some variants with high fluorescence were omitted due to a low percentage of DARPin-displaying cells).

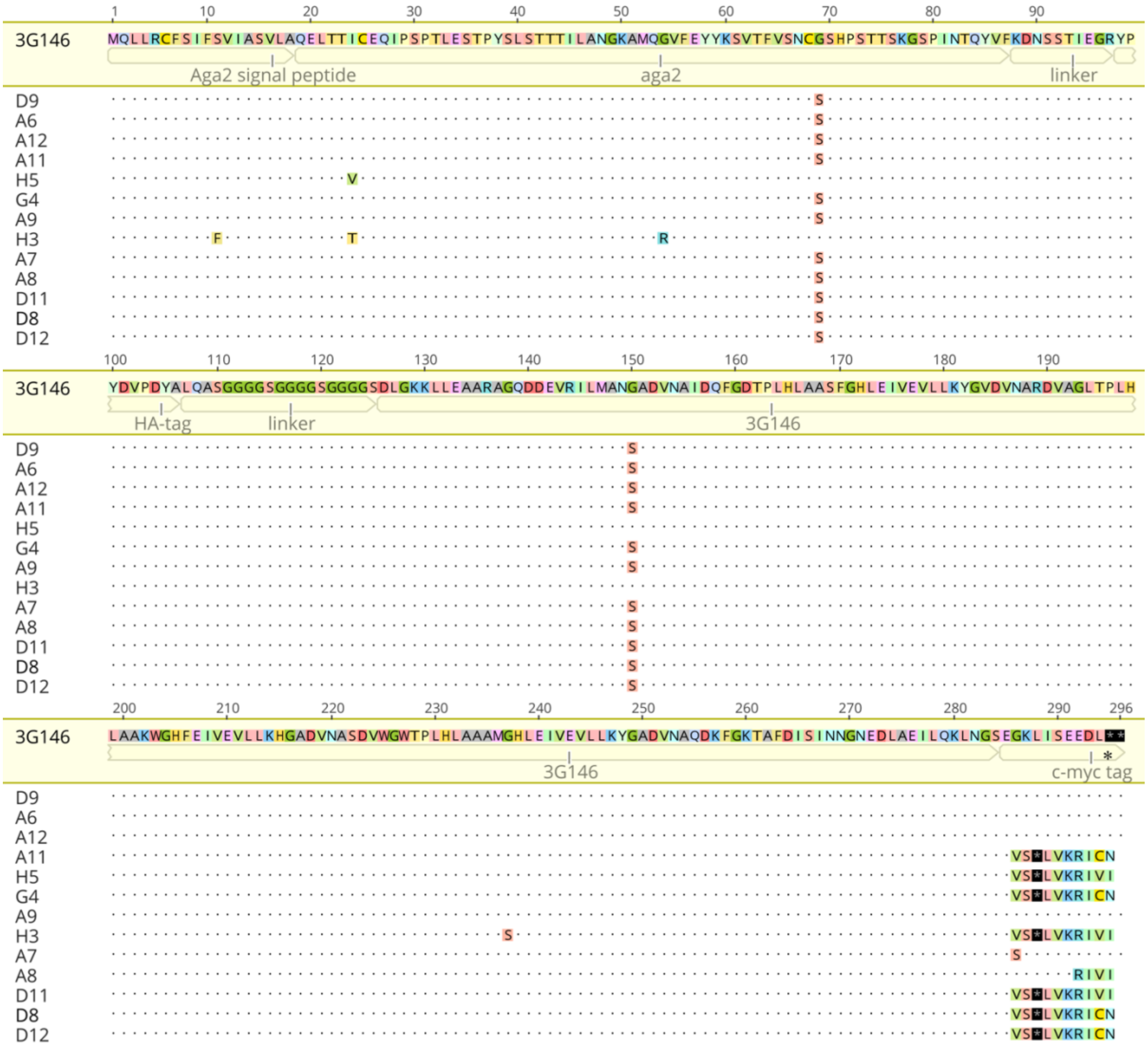

**Supplementary Figure 12. Multiple sequence alignment of variants selected after 2 rounds of directed evolution of 3G146-displaying *S. cerevisiae.*** Wild-type amino acid sequence is indicated in the top row. Individual domains (signal peptide, Aga2, linker, HA tag, DARPin, c-myc tag) are indicated below the wild-type sequence. Variants A6, A9, A12, D9 and A11, D8, D12, G4 have identical genotypes, while variants A7, A8, D11, H3, H5 are unique. Among all variants, two substitutions, G150S and G237S, were found within the coding sequence of the DARPin domain.

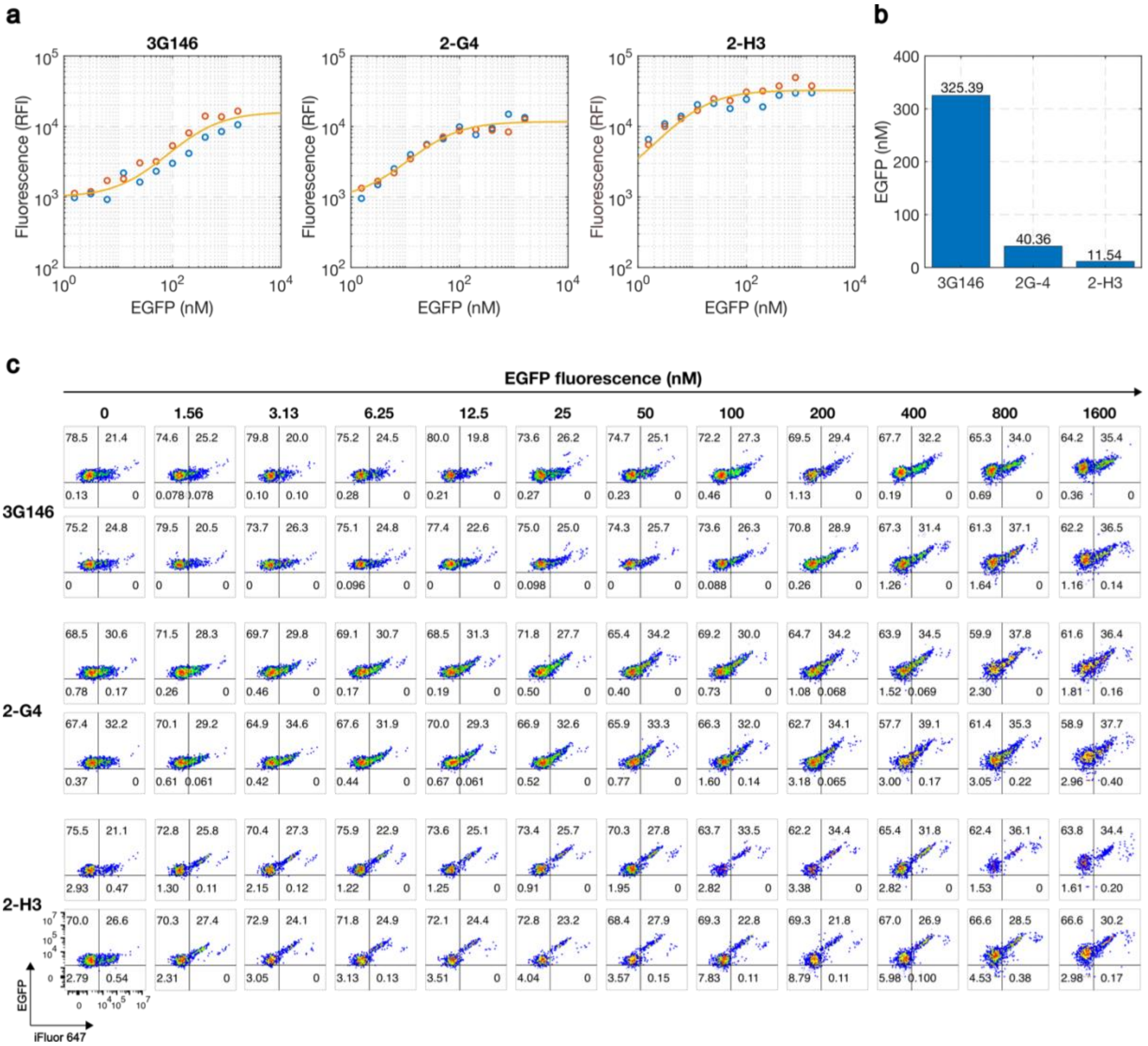

**Supplementary Figure 13. Binding affinity quantification of DARPin variants selected after 2 rounds of directed evolution of 3G146-displaying *S. cerevisiae*, 2-G4 and 2-H3, and wild-type DARPin 3G146 using yeast surface titration.** (**a**) Titration curves showing the background-corrected median EGFP fluorescence intensities of 2 biological replicates of each variant are shown in red and blue. Yellow line represents the monovalent binding isotherm fitted to fluorescence measurements. (**b**) Equilibrium dissociation constants ($K_d$) determined for yeast-displayed DARPin variants. 2-G4 exhibited an 8-fold $K_d$ improvement, while variant 2-H3 showed 28-fold $K_d$ improvement compared to the wild-type DARPin. (**c**) Flow cytometry data (N>1,000) of DARPin variants used for on-yeast binding affinity quantification in **a** and **b**. Variants were cultured in SGΔLW to display 3G146, stained with 1:500 iFluor 647 anti-HA antibody and a range of concentrations of EGFP-His$_6$, and subjected to flow cytometry. Percentage of cells in each quadrant (anti-HA+ EGFP+, anti-HA+, EGFP-, anti-HA- EGFP+, anti-HA- EGFP-) is indicated on the graphs.

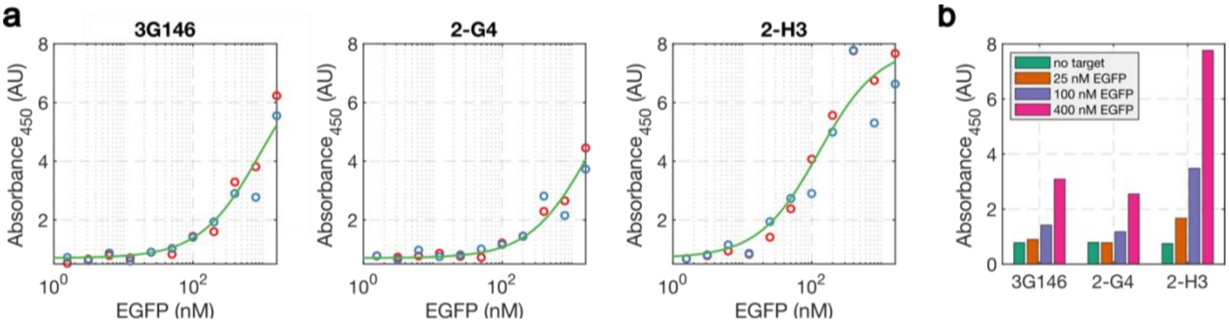

**Supplementary Figure 14. ELISA measurements of purified DARPin variants selected after 2 rounds of directed evolution of 3G146-displaying *S. cerevisiae*, 2-G4 and 2-H3, and wild-type DARPin 3G146.** Sandwich ELISA of serially diluted EGFP target and immobilized DARPin variants. Absorbance values of 2 biological replicates are shown in red and blue. (**b**) Mean absorbance values at selected ligand concentrations – in the presence of 25 nM EGFP, binding signal can only be detected for the variant 2-H3.

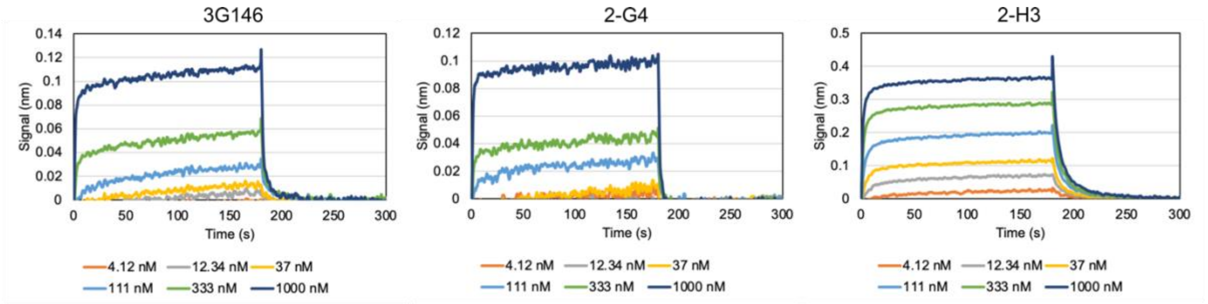

**Supplementary Figure 15. BLI measurements of the interactions between immobilized biotinylated EGFP and DARPin variants selected after 2 rounds of directed evolution of 3G146-displaying *S. cerevisiae*, 2-G4 and 2-H3, or wild-type DARPin 3G146.** Color-coded traces correspond to different DARPin concentrations.

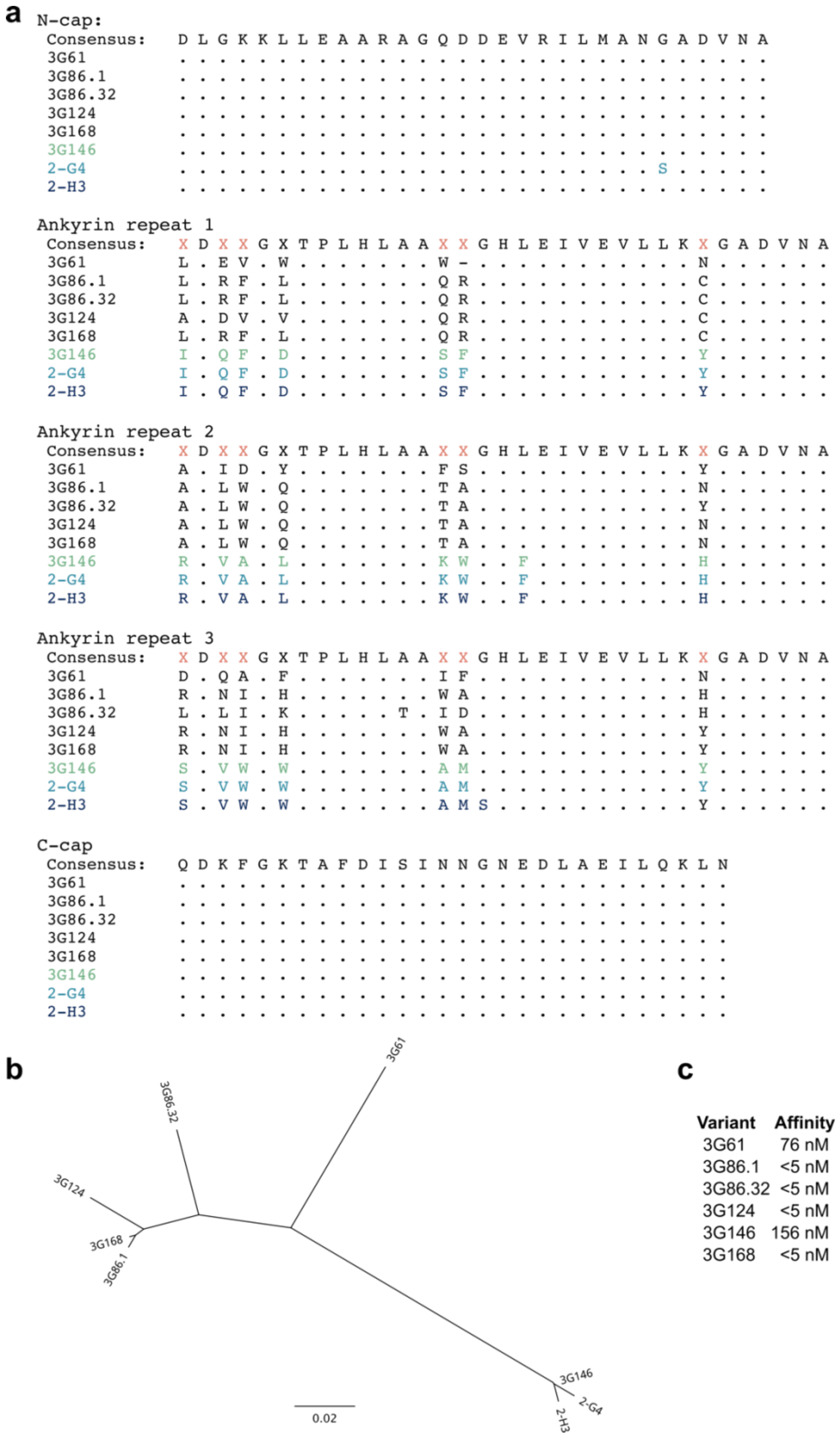

**Supplementary Figure 16. Comparison of variants identified in this study with previously discovered GFP-binding DARPins.** (**a**) Multiple sequence alignment of known GFP-binding DARPins with variants 2-G4 and 2-H3**.** Corresponding DARPin modules are indicated above the sequence. Non-conserved residues that are typically targeted for mutagenesis in DARPin library generation are indicated in red. (**b**) Phylogenetic tree of DARPin variants. Scale bar indicates number of substitutions per site. (**c**) Binding affinity values previously reported for GFP-binding DARPins[13].

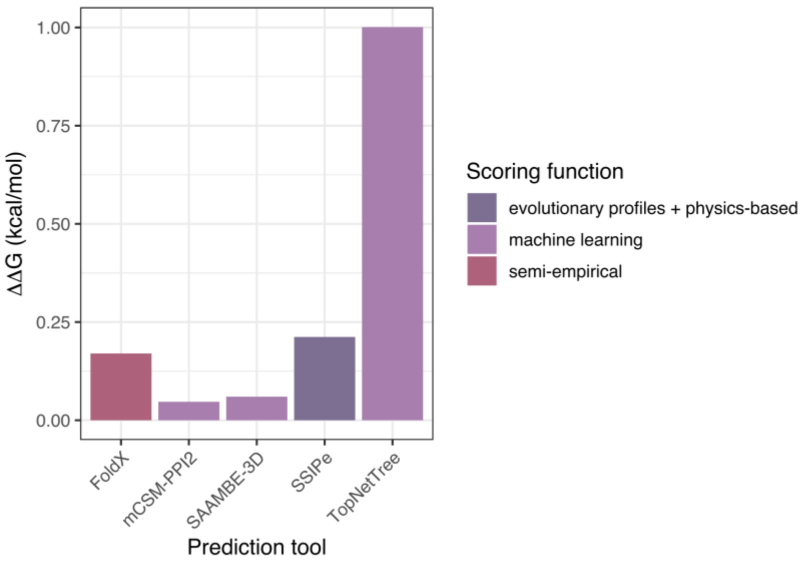

**Supplementary Figure 17.** Predicted binding affinity change for mutation identified in the variant 2-H3. $\Delta\Delta G$ is defined as $\Delta G_{mutant}$ - $\Delta G_{wild\text{-}type}$. Mutations within the range of -0.5 kcal/mol < $\Delta\Delta G$ < 0.5 kcal/mol can be considered as neutral. Tools used for prediction are categorized based on the type of scoring function used to estimate the binding affinity. None of the tools were able to predict the positive effect of the identified mutation on binding affinity with TopNetTree predicting the mutation to destabilize binding, and all other tools predicting the neutral effect.
